# Inter-thinker consistency of language activations during abstract thoughts

**DOI:** 10.1101/546085

**Authors:** Aviva Berkovich-Ohana, Niv Noy, Michal Harel, Edna Furman-Haran, Amos Arieli, Rafael Malach

## Abstract

Human brain imaging typically employs structured and controlled tasks to avoid variable and inconsistent activation patterns. Here we argue against this assumption by showing that an extremely open-ended, high level cognitive task-loosely defined as “abstract thinking” leads to a precise, and highly consistent activation maps. Thus we show that activation maps generated during such cognitive process were precisely located relative to borders of well-known networks such as internal speech, visual and motor imagery. *The activation patterns allowed decoding the thought condition at >95%.* Surprisingly, the activated networks remained the same regardless of changes in thought content. Finally, we found a remarkably consistent activation map across individuals engaged in abstract thinking. The activation to abstract thinking bordered, but strictly avoided visual and motor networks. On the other hand, it partly overlapped with left lateralized language networks. These observations were supported by a quantitative neuronal distance metric analysis. Our results reveal that despite its high level, and varied content nature-abstract thinking activates surprisingly precise and consistent networks in the participants’ brains.

*’Thinker’ refers to: The agent of thought; Intellectual, one who tries to use his or her intellect to work, study, reflect, speculate on, or ask and answer questions with regard to a variety of different ideas” (Wikipedia).*

## Introduction

A prominent experience common to all humans is that thoughts never cease. Experience sampling studies report on 46% of the individual’s awake time spent in thinking ^1, 2^, making it a significant proportion of our cognitive experience. There have been ample attempts to classify thought types, and to suggest their underlying neural mechanisms. One form of thought is abstract reasoning, the capacity to reach novel conclusions on the basis of existing premises - not directly grounded in sensory experience. This process necessarily depends on multiple underlying capacities, but the extent of this reliance on specific mechanisms is a subject of considerable debate ^3^. Given that reasoning and abstraction is of utmost importance to human’s intellectual ability, development and behavior ^4^, a critical question is to understand: What are abstract-thoughts (AT) “made of”?

In using abstract thoughts we aimed to explore the type of reasoning that can be loosely defined as “thinking that encompasses all operations by which cognitive agents link mental content in order to gain new insights or perspectives” ^5^. However, for simplicity of expression, we will exclude the term “reasoning” form the rest of the text and will instead use the word “thinking”.

So far, there were a surprisingly few studies attempting to examine this question. A major hindrance, it appears, is the prevailing assumption that the highly uncontrolled and open-ended nature of such high level tasks will be bound to lead to variable and inconsistent activation patterns. However, as was previously shown ^6^, such an assumption may not be always justified, particularly when the task constitutes natural and prevailing dimensions of human cognition. Here we aimed to examine to what extent the high level nature of abstract thinking, the inevitable changes in content involved and the different individual styles engaged, will lead to highly variable activation patterns.

However, it is important to first clarify what concepts have been proposed regarding to the possible contents of abstract thinking. Several major candidates appear when reviewing the literature: the neural mechanisms supporting language, visual and motor imagery (embodiment), or mentalization. We thereby briefly introduce the theoretical and empirical evidence which supports each one of these views.

### The view that language determines thought

The relationship between thought and speech is one of the oldest yet still debated problems in psychology, logic and linguistics ^7, 8^. Indeed, various linguistics claim that thoughts and speech are strongly inter-related ^9–11^, and that the representations afforded by language are likely to be central to thinking ^12–14^, to the point that language shapes thought and world-view ^11^, albeit other linguistics argue against this view e.g.^15^. Early studies attempting to examine this question through behavior and introspection have revealed mixed results ^8^. Yet, some neural evidence supports the connection, by showing for example that abstract concepts elicit greater activity in the inferior frontal gyrus and middle temporal gyrus compared to concrete concepts, suggesting greater engagement of the verbal system meta-analysis by^15^. In addition, volume reductions in brain areas involved in language have been found to be associated with formal thought disorder in schizophrenia ^17^. Other studies have attempted to map the brain activations associated with inner speech, a potential component of AT, showing different results obtained by different research groups reviewed by^18^.

### Visual imagery and its connection to abstract-thought

Another possibility is that thinking and abstraction proceed via the use of quasi-perceptual or embodied mental models, in which case the high-level spatial and perceptual representations upon which the models are built would be critical for thinking ^19, 20^. This is because our thoughts rely on constructions we build from and meaning we assign to what we sense or recall. A different but related line of research (based on intracranial recordings of neuronal activity conducted in patients) suggests that abstract representations of persons and objects, which are being represented by ‘concept cells’ in the medial temporal lobe, may play a role in building conceptual associations ^21^. A meta-analysis ^16^ showed that concrete concepts (e.g., focusing on the means required to achieve a specific goal - the ‘How’) elicit greater activity in the posterior cingulate, precuneus, fusiform gyrus, and parahippocampal gyrus compared to abstract concepts (e.g. focusing on the purpose of an action - the ‘Why’), suggesting greater engagement of the perceptual system for processing of concrete concepts, likely via mental imagery. Interestingly, a more recent attempt to differentiate the neural activity associated with abstraction and concretization revealed that concretization was associated with activation in frontoparietal regions implicated in goal-directed action, in support of the embodied cognition view, while abstraction was associated with activity within posterior regions implicated in visual perception ^22^.

### The ‘mentalizing network’ – the default mode network (DMN)

Several studies ^23, 24^ have shown that abstraction mindset (thinking ‘Why’) activates the ‘mentalizing network’ – a widespread network of regions associated with theory-of-mind, including the temporal pole, the medial prefrontal cortex, the precuneus and the right temporo-parietal junction. This network largely overlaps the default mode network (DMN), active when we engage in internally oriented rather than externally oriented tasks ^25, 26^. In support of that view, it was shown for example that thoughts arising while viewing artworks are related with DMN activity ^27^.

A potentially informative strategy to assess which of these functions actually dominate abstract thinking can be employed by taking advantage of the recently obtained detailed mapping of the functional specializations in the human cortex. Such knowledge has been accumulated at an unprecedented rate, mainly based on functional brain imaging using functional magnetic resonance (fMRI). Thus, detailed “atlases” that define the functional selectivity of literally every voxel in the human cortex are now available through thousands of mapping studies ^28, 29^. Against this backdrop of detailed known functional atlases, it thus may become feasible to deduce the cognitive components involved in abstract thinking simply by relating the brain activations when participants engage in such abstract thinking processes to the known network selectivity of the cortex, including language, visual imagery, sensory-motor tasks and intrinsically oriented functions. Here we report on the results of an experiment in which such comparison was achieved directly within individual participants.

A second issue of great interest addressed using this direct comparison is whether different contents of thought produce differential activations. Finally, we ask whether individuals that engage in abstract thinking activate similar brain networks, in an analogous fashion to, e.g. sensory activations when confronted with identical stimuli ^5^.

## Results

Our analysis included twenty out of twenty-three healthy participants (see methods) that underwent three fMRI scans (figure 1A): in the first and second scans the abstract thoughts condition (AT) was compared to both visual imagery and language tasks, while the third scan was a Visual Categories scan used to define visual and DMN regions. Subsequently, the participants were interviewed about their experience.

**Figure 1:**
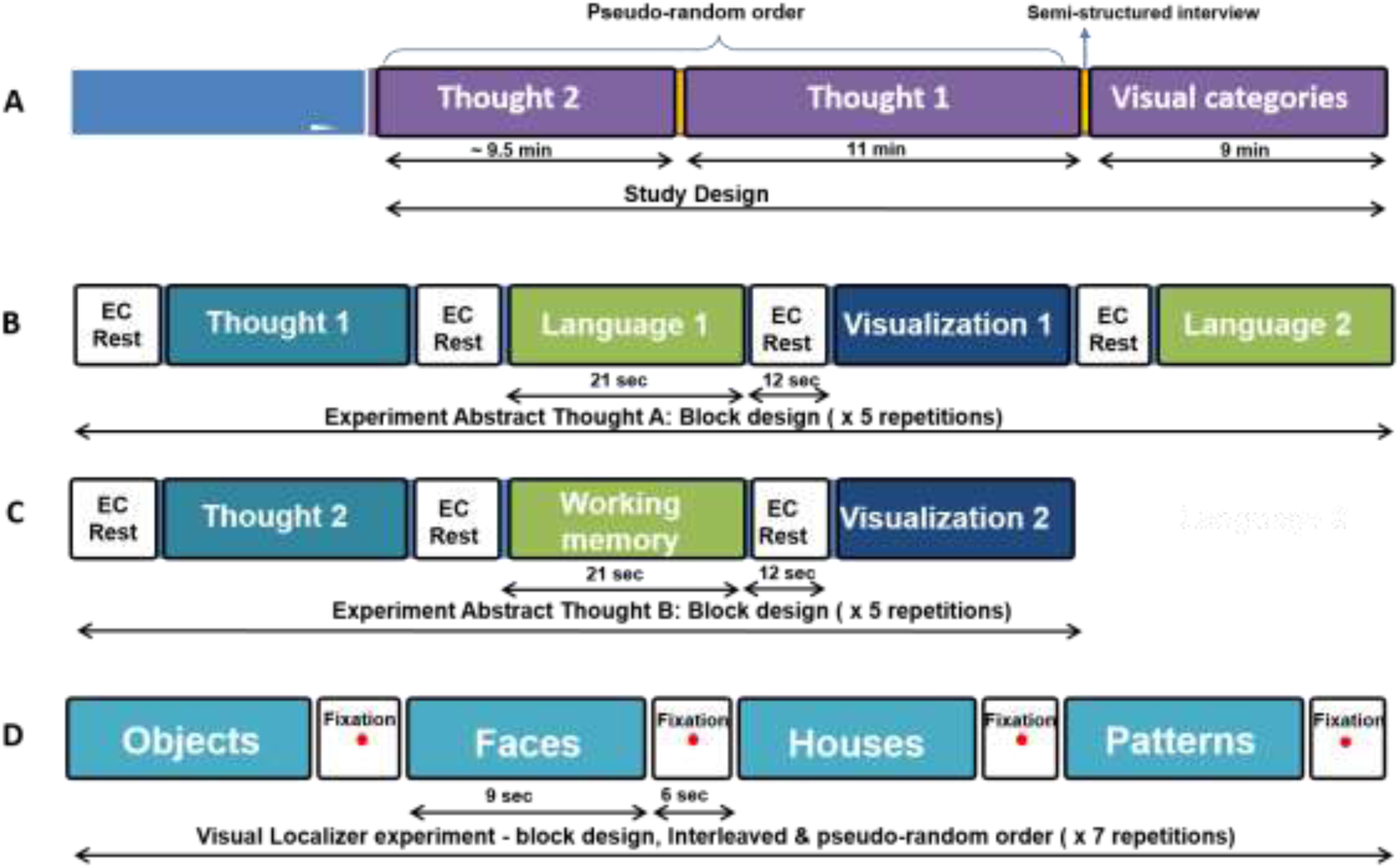
Study design. **A.** The study consisted of three consecutive scans. **B.** Describes the first scan, named “Abstract-thought A”: “EC” denoted eyes closed; “Thought 1” denotes the thought “Is there free will?”; “Language 1” denotes Sentences Conjugation task; “Visualization 1” denotes Imagine Tools task; “Working memory” denotes Working Memory task. **C.** Describes the second scan, named “Abstract-thought B”: “Thought 2” denotes the thought “Is man’s nature good or evil?”; “Language 3” denotes Verbal Fluency task; “Visualization 2” denotes Imagine Navigation task. **D.** Describes the third scan, which includes the Visual Categories task. Purple denotes scans; Turquoise denotes abstract-thought tasks; Green denotes language-related tasks; Dark blue denotes visual-imagery tasks; Light blue denotes visual stimulation tasks.

### First-person reports

The personal reports of the mental content during the different conditions are provided in Table 1. As can be seen, mental contents during the two abstract thought conditions were different for all participants, irrespective of the order of the scans (given in column 2).

**Table 1:** First-person reports. Each participant (n = 23) provided a short personal report of the experience during the abstract thought (AT), rest, and visual-imagery conditions in each of the abstract-thought scans (the order of the scans is given in column 2). * From the reports given by participants 7 and 23 in the abstract thought 1 scan, it could be understood that the contrast AT1 vs. rest could not be representative, due to conceptual mistakes (marked by one asterisk), thus these two participant were excluded from further analyses. **This participant was excluded due to missing neural data.

| Participant | 1<sup>st</sup> scan | Abstract thought 1 scan |  |  | Abstract thought 2 scan |  |  |
| --- | --- | --- | --- | --- | --- | --- | --- |
|  |  | Thought 1 - Is There Free Will? | Rest | Imagine Tools | Thought 2-Is man's nature good or evil? | Rest | Image Navigation |
| 1 | 1 | I thought about what is desire and freedom, afterwards I went into it and I thought about whether it is related to control, and even deeper into it. | I tried to focus visually on the inside of the eyelid in order to not think too much, I saw darkness | In a jewelry making store, I imagined I was holding them | It started very verbally, I analyzed what is bad and good and what is human's nature, then it became more abstract, and in the end I went back to literal meaning and patterns, as I learned to do in sociology | I did the same thing as before, but it was harder to break away from the previous task | From the perspective of levitation, I imagined familiar routes |
| 2 | 2 | The Torah was the starting point and from there I asked questions about religion and democracy | I sang in my heart with thoughts in the background | I was in New York, and pictured kitchenware, and called them by their name | I thought about parashat (section of the torah) "Ki Tavo", blessings and curses, I thought about Noah and wondered why should one be bad or good, what is the profit? | There was a song in my heart, I attempted to continue the song from where I left off last time, with thoughts running in the background | I always walked in the same route in New York |
| 3 | 1 | I tried, there was not enough time. I presented my opinion and arguments with pros and cons. There's this idea that things you're doing come back to you, I was thinking about whether I had free will during this experiment, I was leaning on old thoughts in this subject | I easily disconnected from the task, there was nothing specific | I imagined a housewares shop | I thought about whether the effect was genetic or environmental (nature or nurture), and was it good or bad | I caught myself thinking about my chores at the start of my resting phase | Tracks that I walked on near my house, through the eyes of the walker |
| 4 | 1 | I do not believe in free will because the head is activated by chemistry, for a very long time I think this way | I made an effort to clear my head from any thoughts | I imagined a box of tools | All human qualities are dual, but each quality is dominant at a different certain time | I tried to erase every thought | Through houses, from my house to work |
| 5 | 1 | I tried to build a neurobiological argument and to test it | I felt that the rest disconnected me from being awake and in focus | I went through a variety of tools, I did not succeed in separating the tool from what it is used for | I tried to define it, it was very visual, every time I attempted to develop a discussion, what followed were images | While I rested this time, it was harder to let go of the tasks, especially after the language assignment | I hovered in the air over familiar routes, everything went by very fast |
| 6 | 2 | It was a new thought that I had not encountered before, therefore it was interesting | I had awareness of bodily sensations, both real and those stemming from internal dialogue | It was difficult, I am not familiar with many tools and I started to fill them with content | It translated into a question: is there meaning in the evil projected towards me or others, and is there a choice, do we serve something positive even when we do something evil? | I was calm, the feeling was positive and quiet | I imagined that I was running in a route, like in life |
| 7* | 1 | I forgot the precise instructions and thought of abstract contents related to work * | The body was very relaxed, I did not have many thoughts, rather, I concentrated on my physical sensations | There was a familiar kitchenware tool in the shop, I saw the details | In the beginning I was speaking to myself, and then having a conversation with someone else, like a dialogue | Like before, but it was harder to calm down and return to rest | I walked in familiar and new routes |
| 8 | 1 | I tried to examine the meaning of free will, and the fact that a person lives in a society | Not a lot happened, there was awareness without having to continue the previous condition- that in itself was a thought, I just listened to the magnet and breath | It was difficult to see tools in a static form | I broke the sentence into factors, what is the absolute good and evil? What is human nature - a common thing? Afterward, I asked what was the basis for this - at what stage do we examine this? Before birth? | Quite similar to the former. In the transition to resting phase there is an image of closing a book, connected to closing the previous task | From home to work and reference points on the way, I walked every time from the point I stopped |
| 9 | 2 | It was a complex process for all kinds of reasons, I thought of experiments that involved lifting a hand and decisions and so forth | I had a more difficult time disconnecting from this session in comparison to the previous task | I visited four different stores | I thought about what it means when we do bad things? It does not indicate evil, there is a natural inclination and there is a choice. I had a prior opinion, therefore, it was not challenging | I succeeded in letting go of the previous task | (missing data) |
| 10 | 1 | I thought about how in the experiment I was beginning and stopping the thinking, and whether I could ever stop thinking | I thought about how it is impossible to stop thinking at once, I was wondering what my pupils were doing, and whether it changes from condition to condition | I imagined a warehouse for kayaks, their shape and how I held my hand | I thought about what defines good and evil, is it relative or absolute? How is it defined in different religions? Does a good person define the good in the same way as a bad person? Does the definition changed depending on the definer? | I retracted to thinking about algorithms from work, I felt a physical discomfort from lying down for a while | I walked in the jungle and saw monkeys in the trees, I rowed in the ocean and saw the shoreline |
| 11 | 2 | I thought that desire has a temporal aspect, whether it exists all the time and whether it has meaning when it is not remembered | The assignment kind of slid inside, into my chest | Tools are for motion - it was hard to see them static | I thought about good and evil, what it is for me and another; however, I did not get to the question of the nature of man | The experience was quite physical, part of the time things from the previous blocks returned | I walked home from work; however, I did not feel my body |
| 12 | 1 | At first it was hard to enter the thought because I skipped between topics, in the end it became simple | I tried to follow a method, I always thought about releasing tension from the body | Every time I took a tool and thought about how it worked and why I lacked it in my work | I tried to build a dialogue arguing for and against, I experienced a little excitement from the verbal duel | There was a little inertia, it was hard for me to break away from the previous task | I walked to the preschool from home |
| 13 | 2 | I managed to have new thoughts, and not think of past thoughts in the field | After the language assignment it was harder to break away, I was focused on whether I succeeded | For the most part work tools and houseware tools | I tried to think of a new thought; however, I was having a hard time disconnecting from my previous thoughts because I am a criminal attorney | The resting phase really helped clear my head | Sometimes I walked, sometimes I drove |
| 14 | 1 | For the possible freedom, I thought about situations in which I would do anything to be free | I imagined turning over on my side and falling asleep - every time it took a while to get there, because I was preoccupied with a sense of defeat or success from the previous task | Every time I was with different agricultural tools | This time was more interesting. I thought that we are born virgins and only afterwards adapt | It's hard for me to rest when I wait for the next assignment | A bike trail and driving in a car |
| 15** | 1 | I thought there was free will, I repeated the argument in favor of free will | I had a difficult time with resting after the assignment, it took me a few seconds | Kitchenware in various combinations | I raised arguments both for and against, there was an active discussion | It was easier for me to return to resting in comparison to the previous time | Part of it was in the car or when walking, part of it while hovering |
| 16 | 1 | I discussed the issue of whether the will is truly free or dictated by society and expectations, it was like an internal discussion | I tried to calm down my mind, like a vacuum, not to think about anything. I think I succeeded, without thoughts and feelings | I'm good at it and know the tools | I thought about whether man is born neutral, and that traits are acquired during socializing. In general, man is supposed to be good from his youth. That's the way I depict the Nazi's in my head | I tried to be consistent and to stop the processes during rest, completely, do be completely detached, like a complete shutdown | I saw routes in great detail, like in google maps |
| 17 | 2 | I tried to think about free will, I feel that I succeeded in it | There were fewer changes in the sense of time and less leakage of blocks into the two | Musical instruments and drills | I moved between Hebrew and English, it was hard, but I managed to think about related things, it was associative | It felt as if the resting time was much longer than the assignment, there was a change in the sense of time | I hovered over the passing landscape |
| 18 | 1 | I thought that education affects the will, and the past limits free thinking. I have come to the conclusion that history seems to limit ability, we only want what we imagine | It was cold, I sank into the tone, it was like anesthesia. I felt calm. There were many thoughts | A tool that I need as a draftsman | I thought about people I am familiar with at first, then I analyzed the definitions of good and evil, and thought about our nature to do anything for survival | At first, the transition from the thought to resting was difficult, but with an ambitious attempt, I did succeed in disconnecting | In the beginning while sitting in the car, afterwards it was like one was navigating with a map |
| 19 | 1 | I imagined people arguing, and I brought contradicting thoughts | I imagined a computer screen, throwing something into the recycling bin, so I threw the thought away, and other thoughts entered | I thought of game pieces and bathroom accessories | It was like an argument, a debate between two representatives, claiming opposite arguments | It cut me off from the previous assignment, and then different thoughts entered my mind | I rode a train through familiar routes in between houses |
| 20 | 2 | I made arguments for and against, I thought it could be tested in the MRI | It was hard to get rid of the letter combinations that preceded the rest | In the beginning it was great, and then I finished all the working tools | I thought about whether children are good, how we define good and evil, individual in reference to the general public | I successfully disengaged from the previous task | It was not interesting, but it went well seeing routes |
| 21 | 1 | I thought about what limits our thoughts, what stops us, education? | I treated it as a task, to disengage from the previous task | I went to Holmes Center, I saw tools and focused each time on something different | I did not remember the question, so I thought about what is a thought, is it spirit or matter | I felt a little sleepy during the session, and it was difficult for me to exit my rest for the assignment | I drove in a car between houses |
| 22 | 1 | I thought about free will, and I continued to focus on my previous thoughts | While resting I really did not have any thoughts, I was trying hard for the assignment | Interesting kitchenware | I had a clear opinion, but nevertheless I tried to develop a discussion - but everything went to one side | I tried to automatically analyze what was in the previous stages, and I prepared myself to answer questions | I drove in a car from my home to work |
| 23* | 2 | The topic was less interesting | I thought that the resting phase is associated with the thought, and I continued to think about the nature of man * | I imagined an instrument and a work tool | It was interesting | I saw the sky and the ocean and I rested | I drove in the car and once walked |

To examine the possibility that individual participants relied on different cognitive strategies, and thus belong to different phenomenal types with which might show different neural patters, we used the personal reports to create phenomenal groups. As the free descriptions showed variable content (see Table 1), we used the personal scaling of the verbal and visual content. While the verbal content was always high (graded as 4 or 5 on a 1-5 Likert scale, accept for one participant who graded 3), we used the visual grading, which was more heterogeneous. Participants were divided into to two categories, low visual content (n=12) and high visual content (n=3), chosen as those who graded the visual content as 1 or 2, vs, those who graded 4 or 5, respectively.

### Performance measures

Performance on the Letters WM showed a high (97% ± 7.3) success rate across tested participants. For the two abstract thinking conditions there was no significant self-reported difference between thought 1 and 2 (on a 5-point rating scale): participants rated the level of internal speech as 4.59 ± 0.59 and 4.64 ± 0.58, respectively; the level of internal visualization as 1.82 ± 1.30 and 2.87 ± 1.45, respectively; and the level of interest as 3.60 ± 1.40 and 3.44 ± 0.88, respectively.

### fMRI activations

Turning now to the brain activations as mapped with fMRI (the peak Talairach coordinates of the significant clusters in all the contrasts are reported in supplementary Tables S1-S8), figure 2 shows the average group activation across the 21 participants during abstract thinking about ‘free will’ (AT1-figure 2A) and ‘man’s nature’ (AT2 -figure 2B). As can be seen, both tasks resulted in a robust left lateralized frontal activation, including the supplementary motor area (SMA) and the premotor dorsal (PMD), both in the medial and middle frontal gyrus; Inferior frontal gyrus (IFG) including Broca area (Brodmann area BA-45 and BA-47), as well as middle temporal gyrus (MTG) including (BA-21 bordering Wernicke area), all regions known to play a part in language processing ^30^; AT also activated the temporal pole (TP). Interestingly, despite the different content and subject matter of the thoughts, the activated networks were remarkably similar (see green contours from AT1 map superimposed onto AT2 – figure 2). Moreover, the inter-participant variability was low for the same content (figure 3), indicating that AT produced a robust, highly consistent and specific pattern of activation across different individuals.

**Figure 2:**
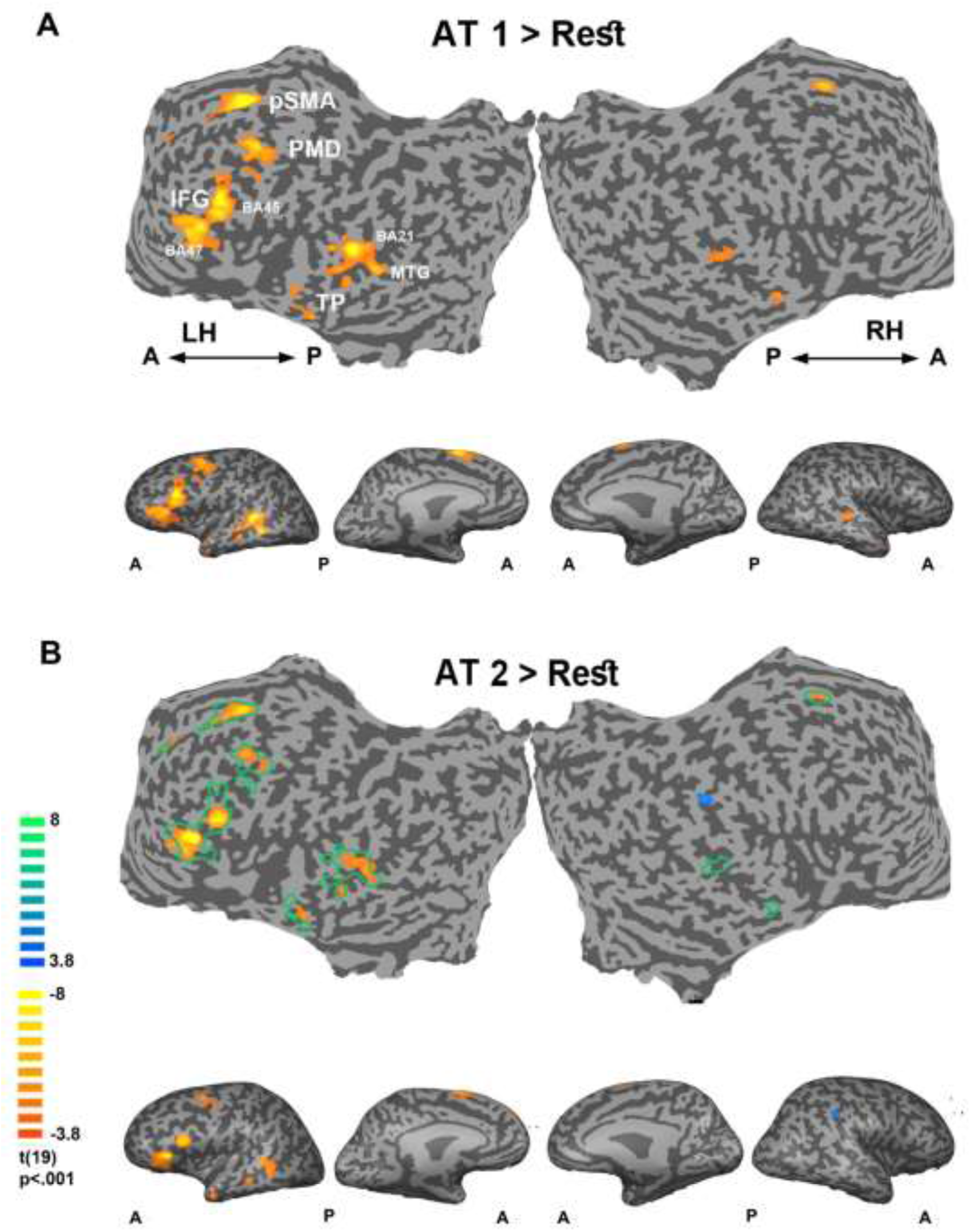
Average group activation during the abstract thought conditions (multi-subject random-effect GLM analysis, *n* = 20, Monte Carlo corrected). **A**. Comparing abstract-thought 1 (AT1 – “is there free will?”) to rest, and **B.** Comparing AT2 (“is man’s nature good or evil?”) to rest. Unfolded (top) and inflated (bottom) views. Color bar indicates activations in yellow, and deactivations in blue. Green contours of the AT1 activations are superimposed on the AT2 activations. Note the remarkable similarity in the activations to the two AT topics. LH - left hemisphere; RH - right hemisphere; A - anterior; P - posterior; pSMA - pre supplementary motor area; PMD - pre motor dorsal; IFG - inferior frontal gyrus; BA - Brodmann area; MTG – middle temporal gyrus; FP – Fronto-polar.

**Figure 3:**
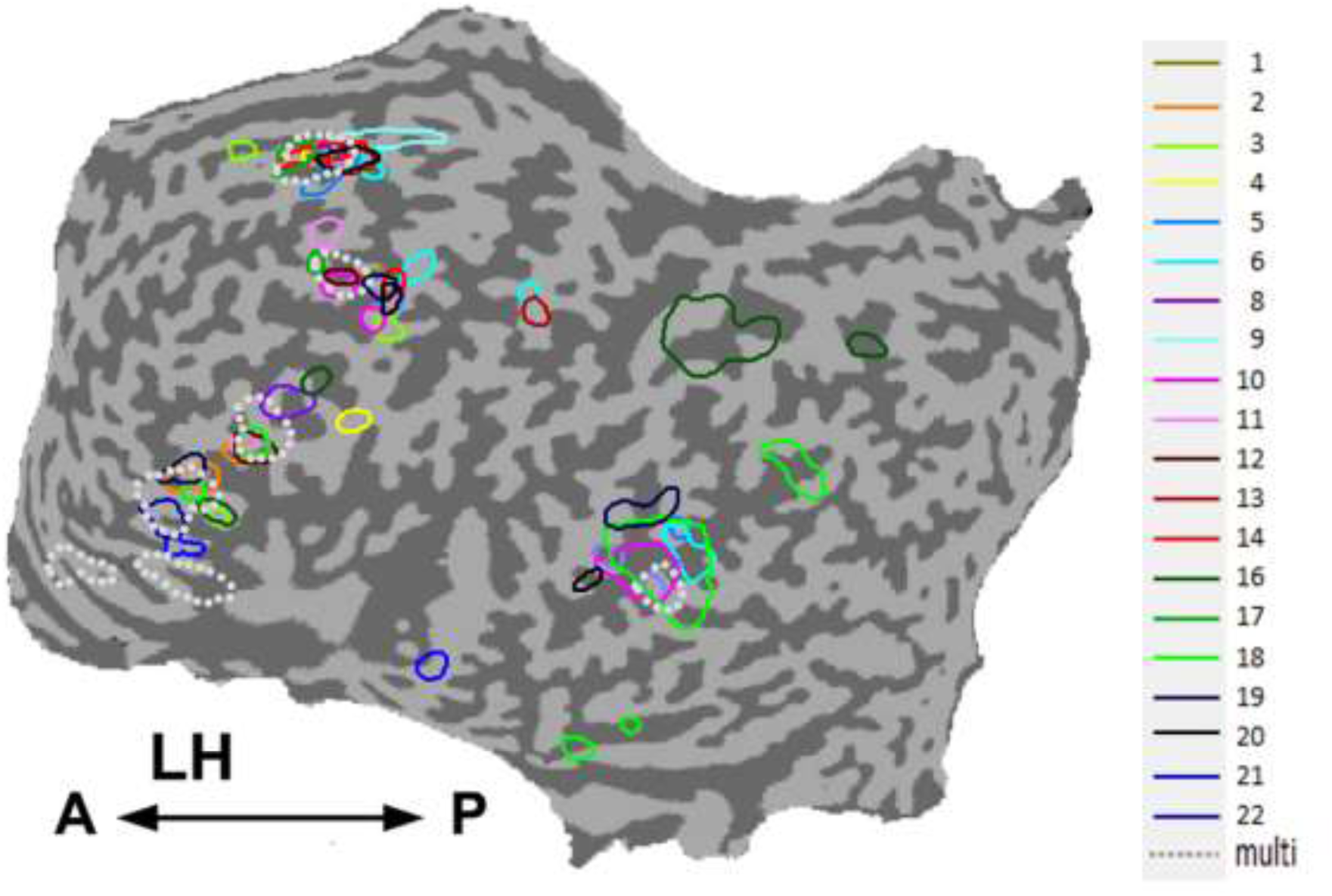
Level of Inter-subject Variability in the left hemisphere in the Abstract Thought 1 task (*n* = 20). The statistical threshold for each participant is > 0.5% activation. Borders of cortical areas were superimposed on unfolded left hemisphere to assess their intersubject variability during the AT1. Colored lines represent boundaries of activated regions for each participant. Note the clustering of activation foci around similar cortical regions. LH, left hemisphere; A, anterior; P, posterior.

To relate the activations during the AT to known cortical functionality at the individual level, each participant underwent a battery of tasks (figure 4) including a Visual Categories scan used to map visual regions and define the deactivated DMN (figure 4B). Conditions of language tasks (figure 4C-E) – were employed to define language-related activations (4D, 3E) as well as working memory activation (4C). Two sensory-motor visual imagery tasks (figure 4F-G) were conducted to allow examination of potential visual imagery components during abstract thinking.

**Figure 4:**
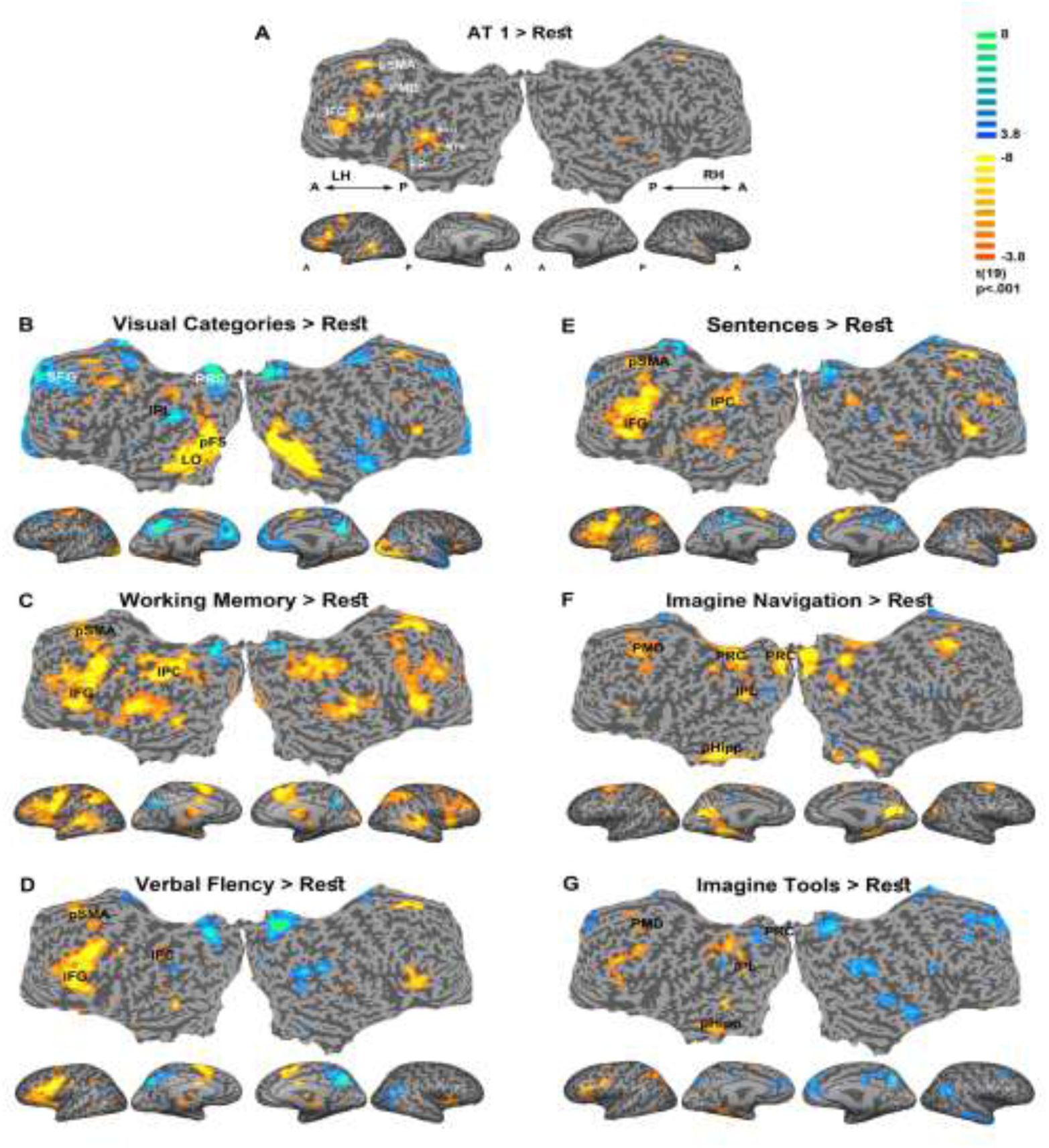
Whole brain maps of activations during the AT 1 and all the other tasks relative to rest (multi-subject random-effect GLM analysis, *n* = 20, Monte Carlo corrected). A. Abstract-thought 1 (“is there free will?”); B. Visual Categories scan: all visual stimuli grouped (four-categories: faces, houses, objects, and patterns); C. Working Memory task; D. Verbal Fluency; E. Sentences Conjugation task; F. Imagine Navigation task; G. Imagine Tools task. Color bar indicates activations in yellow, and deactivations in blue. Note the substantial overlap in activations during the AT condition and the language related ones. Unfolded views: LH - left hemisphere; RH - right hemisphere; A - anterior; P – posterior. pSMA - pre supplementary motor area; PMD - pre motor dorsal; IFG - inferior frontal gyrus; IPC – inferior parietal cortex; SFG – superior frontal gyrus; IPL – inferior parietal lobule; PRC – precuneus; pFs – posterior fusiform; LO – lateral occipital; IPC-inferior parietal cortex; pHipp – para Hippocampus.

In the Visual Categories scan (figure 4B), contrasting all four visual stimuli vs. rest revealed the expected visual-induced increase in BOLD signal in high order visual ROIs, including for example bilateral posterior fusiform (pFs) and lateral occipital (LO) cortices e.g. ^31^. Additionally, it revealed the expected and corresponding task-induced reduction (relative to the fixation baseline) in BOLD signal in the DMN nodes, including the bilateral Precuneus (Prc) and inferior parietal lobule (IPL), as well as superior frontal gyrus (SFG), as previously shown e.g. ^32^.

Subsequently, the BOLD activations in the working memory and two language tasks were examined (figure 4C-E). The Verbal Fluency and Sentence Conjugation tasks showed a typical left-lateralized activation in language regions (figure 4D-E), including inferior frontal gyrus (IFG), Broca area (Brodmann area BA-45 and BA-47), and Wernicke area (BA-21), all regions known to play a part in language processing ^30^. The Working Memory task (figure 4C) showed a bilateral activation generally resembling the other two language task activations. However, as expected, there was a stronger activation at the inferior parietal cortex (IPC, both dorsal and ventral), suggested as the site of the phonological loop ^33^.

The visual imagery tasks (figure 4F-G) showed premotor (PMD) areas, compatible with previous reports ^34, 35^; parahippocampus, possibly reflecting the representation adapted to navigation-related motor tasks ^36^; precuneus (Prc), a key part of the neural substrate of visual imagery occurring in episodic memory retrieval ^37^; and the inferior parietal lobule (IPL), a region related to imitation imagery ^38^ and sense of agency ^39^. The results are in agreement with a previous meta-analysis ^15^ which showed that concrete concepts (focusing on the ‘How’) elicit greater activity in the posterior cingulate, precuneus, fusiform gyrus, and parahippocampal gyrus compared to abstract concepts (focusing on the ‘Why’). Imagery of Tools showed a left-lateralized pattern compared to Navigation imagery, due to the activation of the contra-lateral hemisphere in the right-handed participants during the imagery of manipulating tools, while imagery of whole body movement involved a bilateral activation ^35^. Also, Tools imagery showed a stronger deactivation over DMN regions compared to Navigation imagery.

We then examined to what extent the AT network overlapped any of the cortical activations associated with other tasks (figure 5). This comparison was done by superimposing the activation of AT1 (5A, marked by a green contour) onto all other task-related activations. Comparing BOLD signal in the AT network with high order visual activations (figure 5B), revealed a complete segregation.

**Figure 5:**
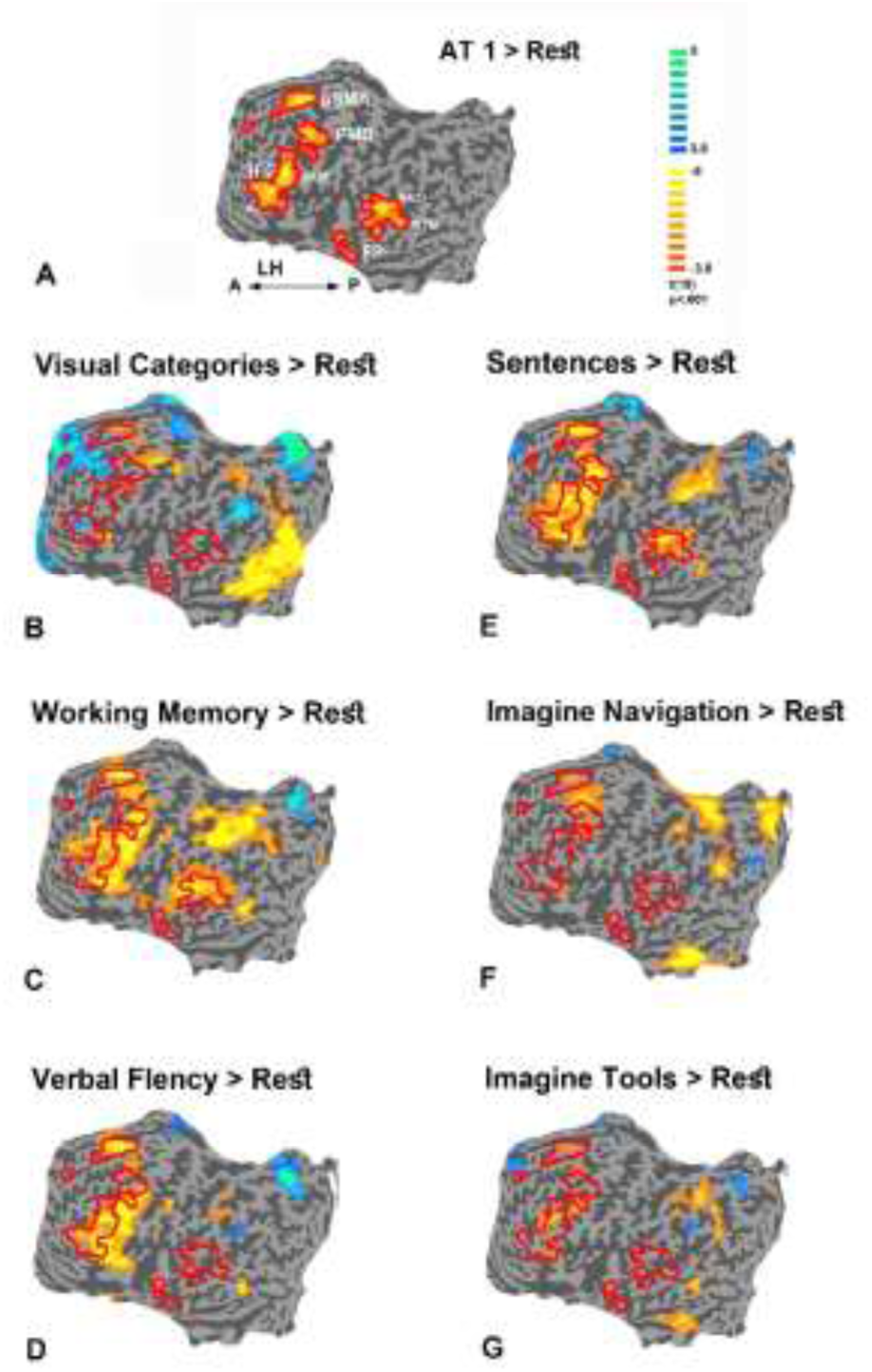
Overlap and segregations of the Abstract thought 1 activations. To examine possible overlaps and segregations of the AT1 activations from the other conditions, the boundaries of AT1 activations were marked by red contours and were superimposed on the other conditions (multi-subject random-effect GLM analysis, *n* = 20, Monte Carlo corrected). A. Abstract-thought 1 vs. rest; B. Visual Categories scan vs. rest; C. Working Memory task vs. rest; D. Verbal Fluency vs. rest; E. Sentences Conjugation task vs. rest; F. Imagine Navigation task vs. rest; G. Imagine Tools task vs. rest. Color bar indicates activations in yellow, and deactivations in blue. Unfolded views: LH - left hemisphere; A - anterior; P – posterior.

Comparing the BOLD activations of the AT network to that produced by the working memory and linguistic tasks (figure 5C-E), revealed a substantial overlap - particularly in more posterior regions (e.g. pre-SMA). However, the AT task activated regions that were not significantly activated during the linguistic tasks. When comparing the AT network to activations generated when the participants engaged in imagery of tools (4F) and navigation (4G) it can be seen that although imagery and AT produced activations in close proximity - there was a complete segregation of the activations for these different tasks.

To study the question of the DMN involvement during AT, we superimposed the deactivation of the DMN during the Visual Categories scan (figure 6A, marked by a red contour) onto the AT and language activations. As can be seen (figure 6B), the DMN was not activated during the AT condition, while deactivated during the working memory and language tasks (6C, 6E, 6F). An ROI analysis on the mean beta values of four DMN regions (Prc and IPL, Bilaterally) reveled that indeed the DMN regions during both AT1 and AT2 were significantly less deactivated compared to the working memory and language tasks (figure 7).

**Figure 6:**
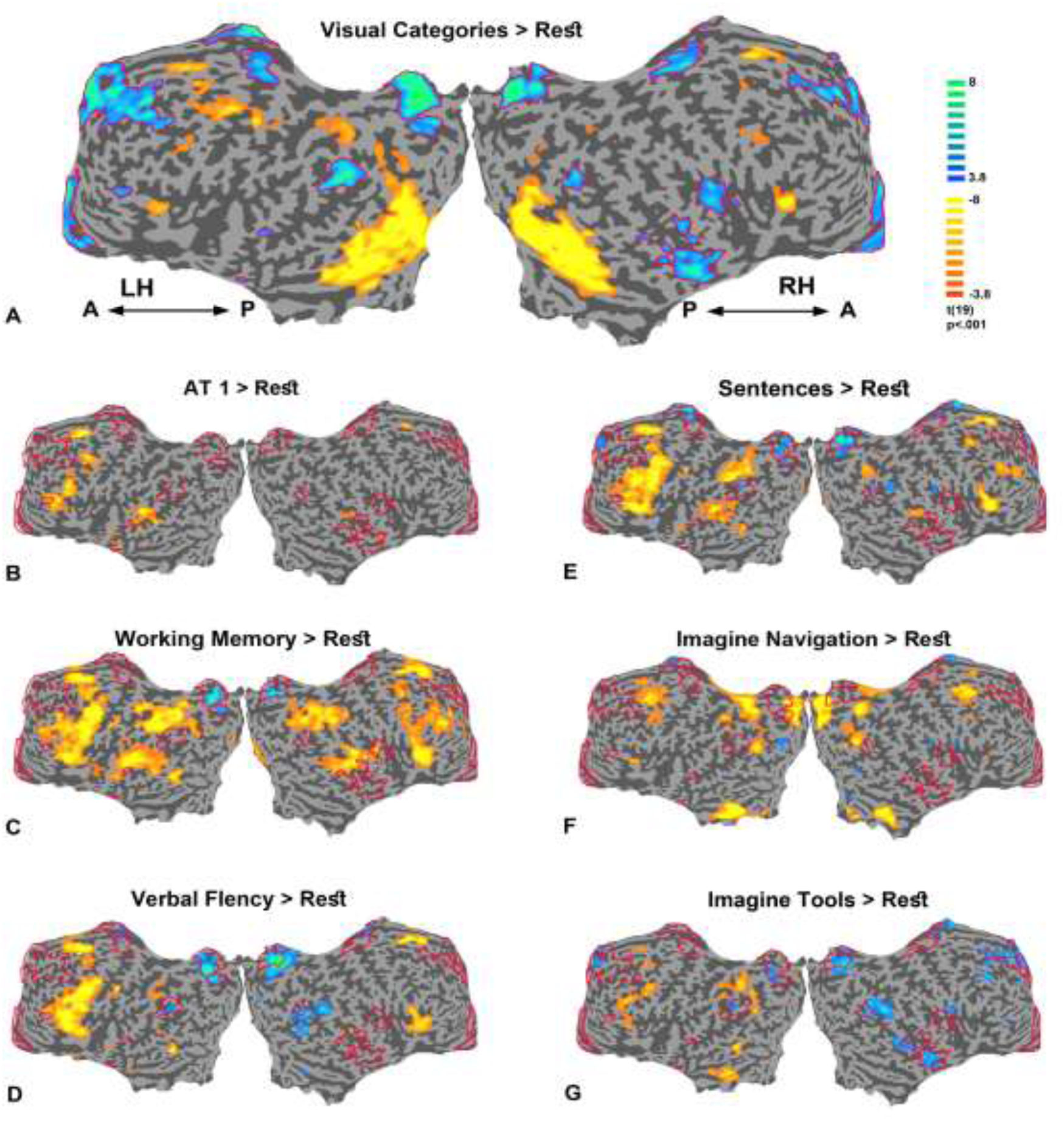
DMN deactivations (marked by red contours) superimposed on the other conditions. A. Visual Categories scan vs. rest; B. Abstract-thought 1 vs. rest; C. Working Memory task vs. rest; D. Verbal Fluency vs. rest; E. Sentences Conjugation task vs. rest; F. Imagine Navigation task vs. rest; G. Imagine Tools task vs. rest. Color bar indicates activations in yellow, and deactivations in blue (multi-subject random-effect GLM analysis, *n* = 20, Monte Carlo corrected). Unfolded views: LH - left hemisphere; RH – right hemisphere; A - anterior; P – posterior.

**Figure 7:**
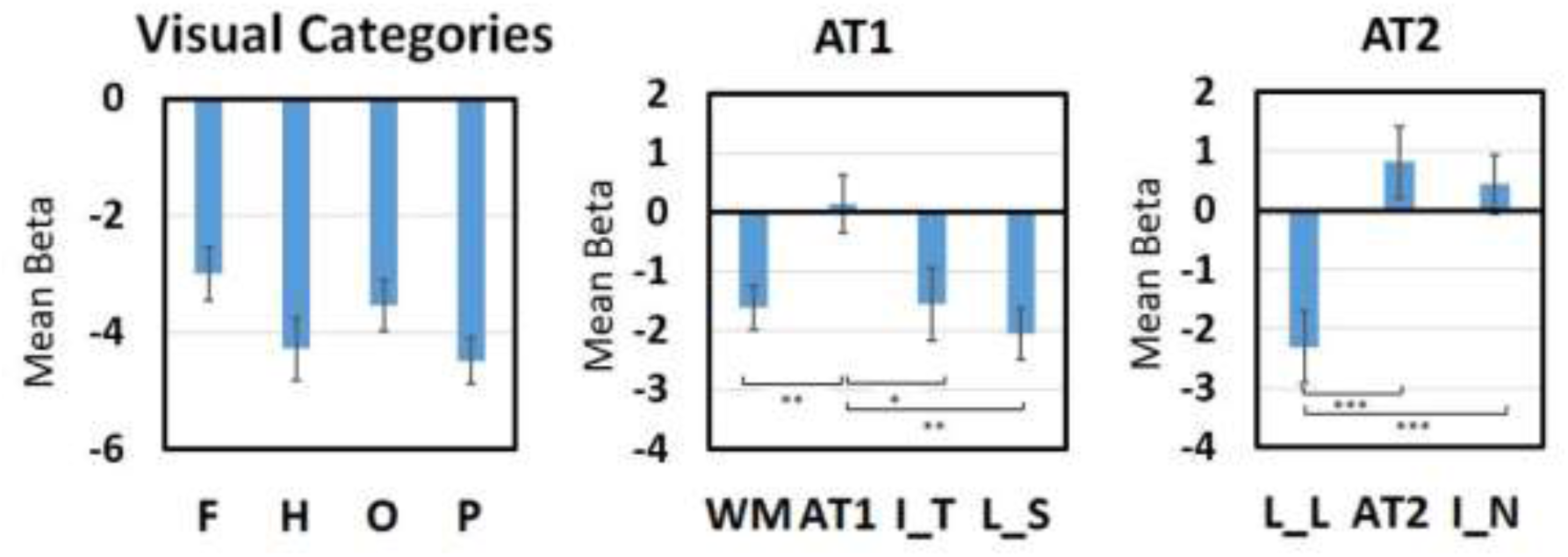
Mean beta values of average DMN ROI. Mean (M ± SEM) beta values (from 4 DMN regions, *n* = 20) are shown for the three scans separately (from left to right: Visual Categories, Abstract-thought 1 and 2). F-face; H – house; O – objects; P – patterns; WM - Working Memory; AT1 – abstract thought 1; I_T - Imagine Tools; L_S – language sentences conjugation; L_L – language letters (Verbal Fluency); AT2-abstract thought 2; I_N - Imagine Navigation. Bonfferoni corrected p values: * p <.05; ** p <.005; *** p <.0005.

An important concern could be that AT were prevalent also during the rest condition. In order to examine this possibility, we compared the AT condition to the rest condition by contrasting both conditions to a third (language) condition. The results are depicted in supplementary figure S1. As can be seen the AT1 and AT2 conditions produced drastically different activation maps when compared to the rest condition. These differences were mainly and consistently evident in frontal regions (see arrows).

### Neural pattern classifier

If indeed the activation patterns of the thought conditions were highly reproducible across individuals, this may allow to successfully predict the cognitive state of the individual participants directly from their activity patterns. To examine this possibility, we performed a multi-variate, leave-one out, classifier approach (see methods for details) in which the pattern of activation of individual participants was compared to the average patterns across the rest of the group for five bulk conditions. When including all cortical voxels, our ability to predict the thought condition (averaged across both thought tasks) compared to the other bulk condition categories (language, imagery, working memory and visual) using this analysis resulted in a remarkable 95% performance (19 out of 20 individuals, chance performance 20%). When including only 50% of the voxels showing low variability the performance increased to 100%. The decoding efficiency of the AT condition, compared to all bulk conditions was highly significant (p<0.05, permutation test, see methods for details).

### Neuronal distance results

The multi-variate approach allowed us also to compare the similarity (neuronal distance - see methods) in activation patterns across the different conditions. The results are depicted in figure 8A-C and supplementary figure S2. Statistical analysis of the bulk conditions revealed that the language tasks were significantly more similar to the AT conditions compared to the other tasks (figure 8C).

**Figure 8:**
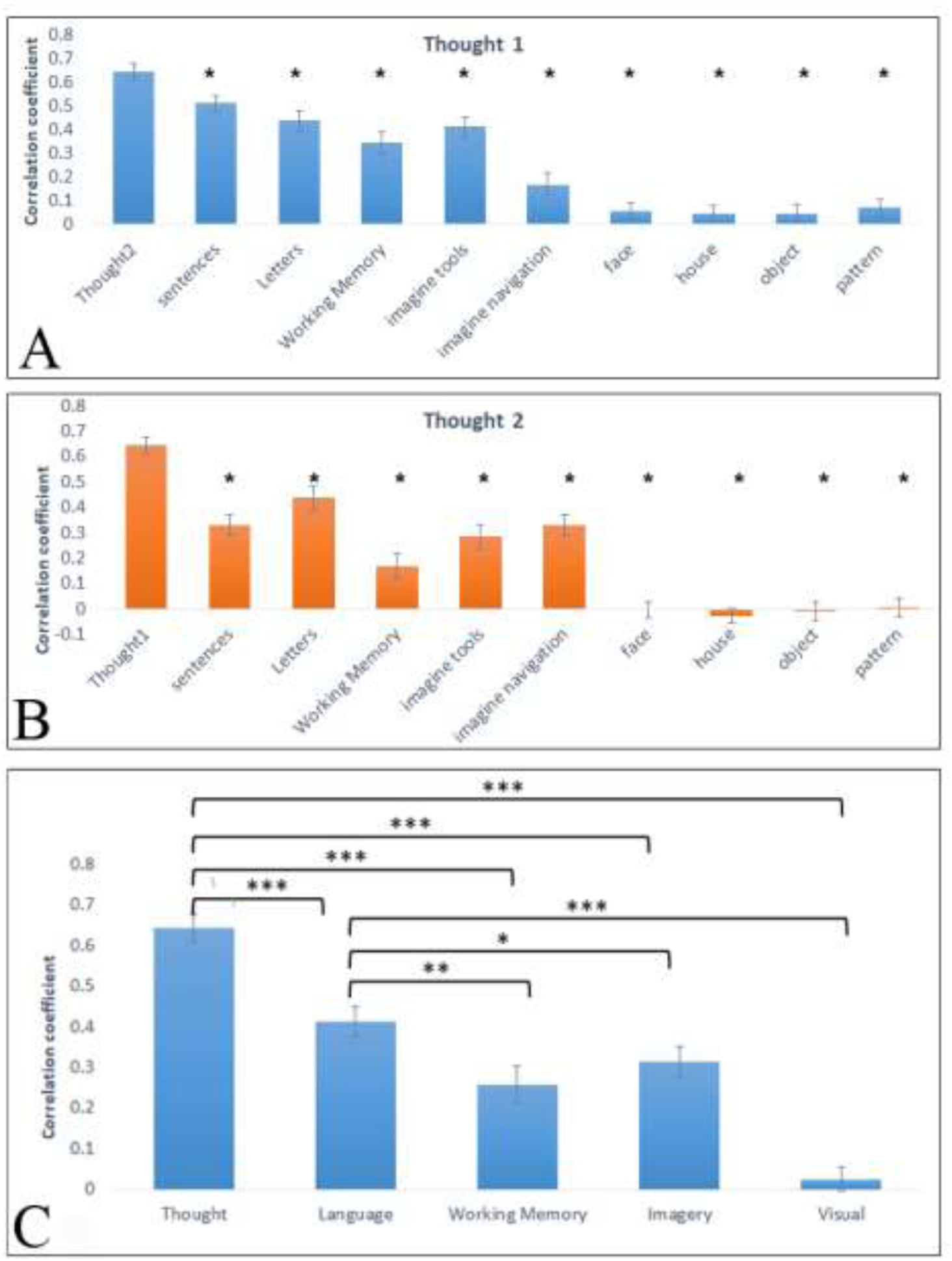
Neuronal Distance Analyses as bar histograms. Mean ± SEM of correlation coefficients (n = 20) between all the conditions and A. AT1 (Figure 2A); and B. AT2 (Figure 2B). C. To reduce multiple comparisons, we lumped together the results of the two thought conditions, the two language conditions, the two imagery conditions and the four visual conditions. * p < 0.05; ** p < 0.0005; *** p < 0.00005.

To examine the possibility that different phenomenal categories might show different neural patterns during the AT conditions, we contrasted the AT phenomenal groups described above. Correlation coefficient between AT1, AT2 and bulk AT was performed with other conditions in each participant. Student’s t-test was performed between the correlation coefficients comparing the two phenomenal categories. No significant difference was found for either AT1 or AT2 (supplementary figures S3A and S3B, respectively). Additionally, average correlation coefficients of AT1 and AT2 with other conditions in the two categories as well as bulk AT with other conditions in the two categories are displayed (supplementary figure S3C).

Similar to our findings in the whole group of participants, AT1 and AT2 had the highest correlations with significant differences found in almost all comparisons to other conditions performed separately in the high and low visual content groups. Exceptions were correlations in the high visual content group, with no differences between correlations of AT1-AT2 compared to AT1-sentences, AT2-letters, AT2-imagine navigation, and in the bulk with AT2-working memory.

### Inter-participant variability

To obtain a quantitative assessment of the inter-participant variability we calculated the distance of each participants’ patterns from the mean for each conditions. The results (average ± SEM) are displayed in bars histogram (figure 9). As can be seen the AT conditions showed a similar levels of inter-participant similarity to all other conditions.

**Figure 9:**
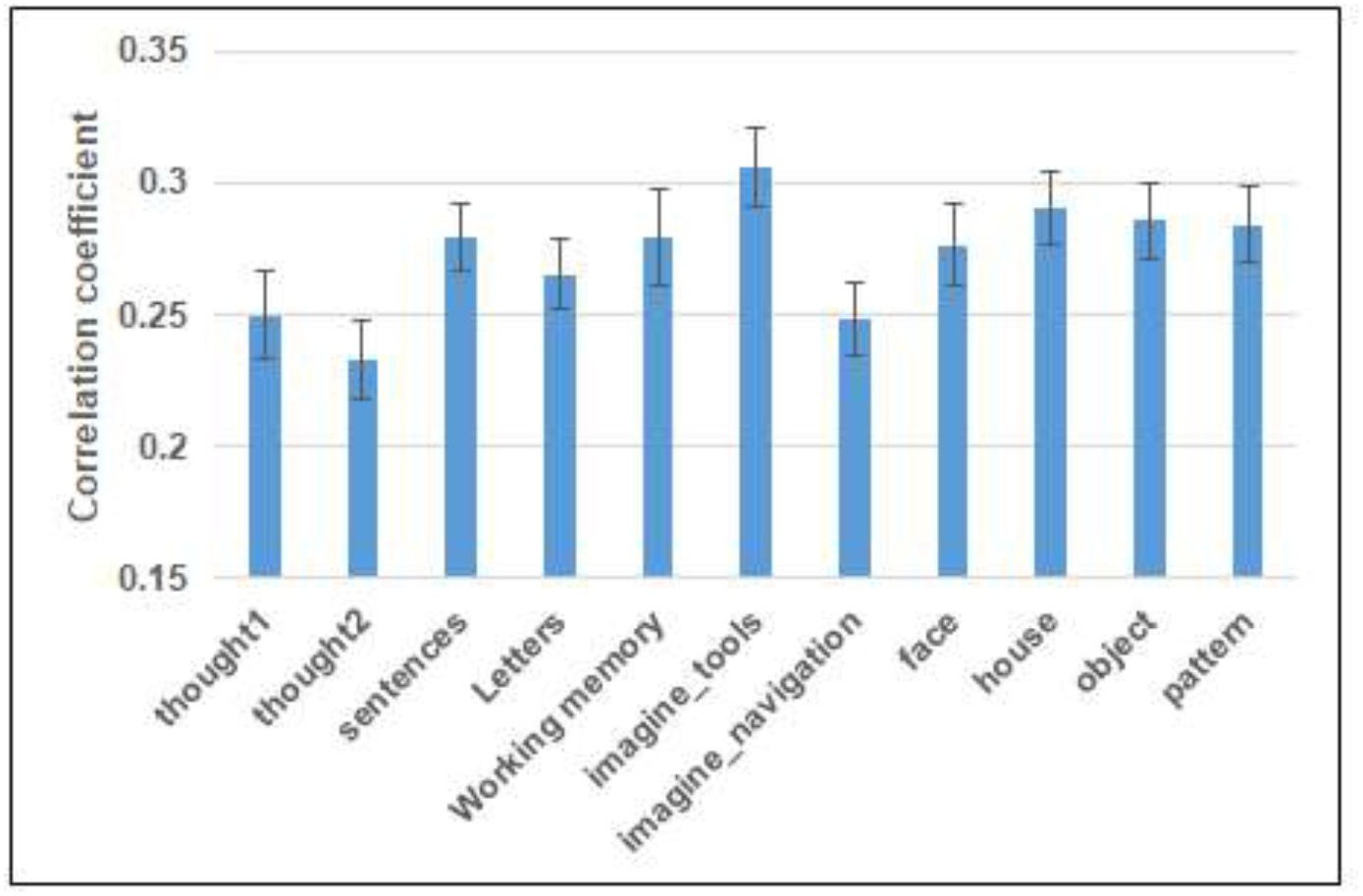
Average (Mean ± SEM) of Correlation Coefficient in each condition between participants (n = 20).

## Discussion

The first issue that our results address is how sensitive was the AT activation to thought content. Our study reveals that the pattern of activation was invariant to the specific content of the AT as illustrated by the remarkably precise overlap between the activations produced by the two types of AT (figure 2). This suggests that different AT contents engage similar networks when viewed at the resolution of BOLD fMRI. The highly reproducible nature of the activation patterns during the AT conditions across individual participants allowed us to successfully predict the engagement of participants in thinking as compared to the other tasks based on their activation patterns alone. This was highly successful when averaging the groups of conditions together (95% predictability for all voxels, 100% predictability for the 50% least variable voxels). Thus, the ability to predict the AT condition from the patterns of activity compared to all other conditions was highly significant, again illustrating the consistent nature of the AT activation pattern across individual participants. It is important to note that the similarity that we observed between the two AT tasks was not only evident qualitatively at the level of univariate responses, but was also significantly higher in our multi-variate pattern similarity analysis (see figures 8 and S1). Furthermore, this similarity was revealed under two independent scans, which may have introduced some mismatches unrelated to the neuronal activations.

An important question concerns the source of the similar activation patterns across the two AT conditions. One possibility could be that the similar activations were generated by a large overlap in the specific contents of thoughts despite the different questions posed for the two AT conditions. The other possibility could be that the activation patterns, at least at the fMRI resolution, could not differentiate between the subtle changes in content-and reflected more general aspects-such as cognitive strategies, recursive thinking ^40^. The similar activations to different AT contents supports the latter conclusion. This conclusion was strengthened by the participants’ reports indicating that their mental content during the two abstract thought conditions were different in all cases (Table 1).

The second issue that our results address is how sensitive were the AT activations to individual differences. We found that despite their high cognitive level and rather abstract and vaguely defined nature, AT produced a surprisingly consistent and specific pattern of activation across different individuals (figure 3). It could be argued that averaging the patterns of activations across a large number of participants may have masked distinct thought contents and strategies in individual groups. To examine this possibility, we searched in the post-hoc subjective reports for indications of distinct sub-grouping of mental content during the AT. We indeed identified one such distinction related to the level of visual imagery during one or both AT conditions. A separate analysis of these two subgroups revealed no significant group differences (supplementary figure S3). This remarkable consistency across subjects was further supported by a quantitative estimate of the inter-participant variability in activation patterns (figure 9) which revealed largely similar levels across the different conditions including the visual stimulation ones. This finding suggests that across individuals the overall cognitive strategies employed during abstract thinking were largely similar, despite the inherent open nature of AT. This surprising similarity is compatible with previous observation in other systems, indicating that participants show a remarkable level of “synchronization” in their cortical activations when confronted with naturalistic stimuli ^29^.

The third issue we address is - what is AT “made of”? – i.e. with which known task-related networks AT activations overlap most prominently? Our approach directly tested several major candidates: visual imagery, mentalization, and the neural mechanisms supporting language. We also compared the AT activations to known task and sensory networks - most importantly somatosensory and motor representations.

When examining the relationship of activations during AT to various working memory and language production tasks, we found a substantial overlap (see figure 5C-E) in language areas (e.g. Broca, Wenicke). This overlap is in line with theories proposing a close association between language and thought. A particularly interesting theoretical link of this kind has been recently proposed by Berwick and colleagues ^41^, suggesting that the emergence of inner thoughts actually set the evolutionary stage for language development.

Additionally, we found a substantial overlap with the language tasks in the supplementary motor area (SMA) and the premotor dorsal (PMD) both key structures for preparation and initiation of a voluntary action ^42^, recently shown to be involved in the sense of agency - the feeling of controlling events through one’s actions ^43^. Such activations were evident in all self-generated verbal tasks employed in our study as well, such as word fluency and working memory (see figure 4C-E). These results are compatible with recent empiric evidence that inner speech - a purely mental action - is associated with an efference copy, previously associated only to overt movements, thus suggesting that inner speech may reflect a special type of overt speech ^44^.

When examining the relationship of deactivations during AT in DMN regions compared to rest, we failed to find significant modulations compared to rest - either deactivations (figures 6 & 7), or activations. This argues against the view that abstract thinking is strongly based on enhanced activation of processes typically associated with the DMN, such as self-related cognition ^24, 45^, “metallization” ^22, 23^ or introspection ^32^.

It could be argued that comparing the AT condition to rest may have under-estimated the brain activation during AT since similar cognitive processes could have been generated during rest. In order to examine this possibility, we directly compared the rest and AT conditions by contrasting them with a language condition. Our results (figure S1) reveal major differences between the AT condition and rest, with higher activation during AT in frontal regions. Thus, although we cannot rule out the possible engagement of some AT during rest, the resulting brain activation during the two conditions was substantially different.

Notably, AT also activated the temporal pole which was not activated by any other task we tested. The temporal pole is suggested to be activated in integrating complex, highly processed perceptual inputs to visceral emotional responses ^46^. This activation appears compatible with the abstract and high level cognitive processing involved in AT.

A recent fMRI study of spontaneously arising thoughts ^48^ showed a set of regions similar to the AT network shown here, including the SMA, posterior cingulate cortex, IPL, temporal pole and superior temporal gyrus. This overlap may be attributed to the rather unconstrained nature of the AT tasks in the present study.

While examining the cortical networks associated with AT activation is important to revealing the cognitive components engaged during such thoughts, not less informative is examining those networks that were not activated during AT.

Comparing the AT activations to high order visual activation didn’t show any overlap (figure 5A). Furthermore, the activation associated with visual imagery was clearly segregated from the AT activations (see figure 5F-G). This result therefore argues against a role for high order visual regions and visual imagery in AT.

Another important cortical region that wasn’t activated, was the posterior parietal cortex (PPC), known to be involved in many basic operations—such as spatial attention ^49^, mathematical cognition, working memory, long-term memory, coordinate transformation ^50^, and language reviewed by ^3^. However, given that evidence suggests that the PPC’s contribution to thinking may be most closely related to its role in mathematical cognition rather than linguistic processing or spatial relations ^3, 51^, it is plausible that the abstract thoughts evoked in the present study did not activate such a probabilistic computation dependent on numerical processing mechanisms.

Of particular interest is the complete absence of enhanced activation in the parietal lobe areas of body representations e.g. ^34^. Such failure argues against a straight-forward embodied cognition view, that the body functions as a constituent of high order cognitive processes ^52^. Of particular relevance here is the suggestion that somatosensory and motor cortices activation constitute an integral part of thought processes ^53^. It appears that at least for the kind of thought processes employed in abstract thinking, such somatosensory involvement was not required.

Furthermore, comparing the level of similarity of each group of conditions to the AT with the neural distance analysis revealed a significantly higher similarity of the language tasks to the AT conditions (figure 8C). Thus, our quantitative analysis further supports the preferential overlap of AT to language-related functionality. However, it is important to emphasize that the activation patterns to AT showed a significant difference from purely language tasks (figure 8C).

To summarize, our results suggest that there is an abstract-thought left-lateralized network, involving regions relevant to language processing, volition, and likely intrinsic processing. Importantly, this network does not include somato-motor representations, or default network activation. Finally, this network is invariant to thought content and highly consistent across individuals.

## Methods

### Participants

Our study included twenty-three healthy participants (age 40 ± 9.5 years, 8 females) that underwent fMRI scanning. Pre-established inclusion criteria include right handedness by self-report, demographic variables (age, education level), fMRI safety criteria, and no history of neurological or mental health disorders (including no medication use). All participants provided written informed consent for their participation. Two participants did not fully comply with the task (see first-person reports in Table 1, marked by asterisks), due to conceptual mistakes: one forgot the precise AT1 instructions (#7), and the other thought that the resting epoch is a continuation of AT1 (#23). Thus, these two participant were excluded from further analyses. In addition, one participant (#15) had missing neural data. Hence, only 20 participants are reported throughout the study, and all figures were adjusted accordingly. The sample size is well above the optimal number of participants suggested for fMRI studies, of about 16 ^54^. The experimental procedures were approved by the Ichilov hospital ethics committee.

### Imaging setup

Images were acquired on a 3 Tesla Trio Magnetom Siemens scanner, at the Weizmann Institute of Science, Rehovot, Israel. Functional T2*-weighted images were obtained with gradient echo planar imaging (EPI) sequence (TR=3000 ms, TE=30 ms, flip angle=90°, FOV 240 mm, matrix size 80×80, scanned volume—46 axial slices tilted to the ACPC plane, of 3×3×3 mm voxels to cover the whole brain without gaps).

The participant’s head was placed on a foam cushion for stabilization, and MR compatible earphones (MR confon, Magdeburg, Germany) that substantially reduce external noise were placed on the ears. Three-dimensional T1-weighted anatomical images were acquired to facilitate the incorporation of the functional data into the 3D Talairach space (1×1×1 mm resolution, 3D MP-RAGE sequence, TR=2300 ms, TE=2.98 ms, TI=900 ms/ 9° flip angle).

### fMRI stimuli and experimental design

The study consisted of three consecutive scans of approximately ten minutes in length (figure 1A, Table 2). Prior to each scan, participants received detailed instructions regarding the various tasks at each condition and were told which auditory cue signified each condition. Presentation^®^ software (Neurobehavioral Systems, Inc.) was used to deliver the auditory stimuli via the MR-compatible headphones.

**Table 2:** Details of scans and tasks used in this study.

| Scan | Condition | Task | Length | Instruction | Cue |
| --- | --- | --- | --- | --- | --- |
| Thought 1 | Abstract thought | Abstract thought 1 (AT1) | 5 blocks (each 21 sec) separated by rest blocks | Think about the question "Is there free will?" | Cue: "thought", cue duration: 850 ms |
|  | Visual imagery | Imagine tools | 5 blocks (each 21 sec) separated by rest blocks | Visualize tools of your preference (e.g. working tools or kitchenware) as you imaginatively walk through a big store and screen the shelves | Cue words: "imagine tools", cue duration: 1250 ms |
|  | Language | Sentences Conjugation | 5 blocks (each 21 sec) separated by rest blocks | Participants were given sentences, and instructed to conjugate them to different tenses/ objects. | "He went to the sea"; "She is buying bread"; "You are going up"; "I will sit in the garden"; "They wore a coat" (respectively: 1550, 1550, 1200, 1800, 1500 ms) |
|  |  | Letters Working Memory | 5 blocks (each 21 sec) separated by rest blocks | Press "1" if the last letter was one of the six letters, and "2" if not | 1<sup>st</sup> cue: each time a different series of six Hebrew letters (e.g. "pe, reish, kuf, gimel, lamed" (cue durations 2700-3200 ms)<br>2<sup>nd</sup> cue: one Hebrew letter |
|  | Rest |  | Each 12 sec | Rest as best as you can | Cue: "rest", cue duration: 620 ms |
| Thought 2 | Abstract thought | Abstract thought 2 (AT2) | 5 blocks (each 21 sec) separated by rest blocks | Think about the question "Is man's nature good or evil?" | Cue word: "thought", cue duration: 850 ms |
|  | Visual imagery | Imagine navigation | 5 blocks (each 21 sec) separated by rest blocks | Visualize a navigation of your route home | Cue words: "imagine navigation", cue duration: 1350 ms |
|  | Language | Verbal fluency | 5 blocks (each 21 sec) separated by rest blocks | Generate continuously and silently new words which start with the cue letter | Cue letters were: "a/b/g/r/n", cue duration: 500, 350, 650, 500, and 730 ms respectively |
|  | Rest |  | Each 12 sec | Rest as best as you can | cue: "rest", cue duration: 620 ms |
| Visual Localizer | Visual | Faces | Each condition had 7 blocks (pseudo-random order), each lasted 9 sec, (followed by rest) - 9 images in each block; (each for 800 ms followed by a 200 ms blank screen) | Fixate on the central fixation dot, and report whether the presented stimulus was identical to the previous stimulus or not, by pressing the buttons "2" or "1", respectively | Stationary gray scale line drawings of faces, buildings, common man-made objects, and geometric patterns (16 of each type). A central red fixation point was present throughout the experiment, including the fixation blocks |
|  |  | Houses |  |  |  |
|  |  | Objects |  |  |  |
|  |  | Patterns |  |  |  |
|  | Rest |  | 6 sec blank screen | Fixate on the central fixation dot | A central red fixation point |

In the first and second scans (Thought 1 and 2), conducted with closed eyes, abstract thought (AT) was compared to both visual imagery, working memory and language tasks. These two scans were given in random order (15 performed first “Thought 1” and 8 performed first “Thought 2”, see Table 1). The first scan (figure 1B) included 4 tasks: i) AT1 task (“Is there free will?”), ii) a classical language task ‘Sentences Conjugation’, iii) a working memory task examining working memory engagement, and iv) a visual imagery condition (“imagine tools”).

The second scan (figure 1C) included 3 tasks: i) the AT2 task (“Is man’s nature good or evil?”), ii) a classical language task (verbal fluency), and iii) a visual imagery task (“imagine navigation”).

Finally, in the last scan (figure 1D), each participant performed with their eyes open a Visual Categories, which was used to define the DMN regions for our group of participants, as previously reported ^55^.

Immediately following each of the first three scans, each participant was interviewed while in the scanner via an intercom with a semi structured interview, in order to provide a brief description of the content of their experience. Particularly, participants were asked after the first and second scans to report feely the mental content during the abstract thought (AT), rest, and visual-imagery conditions – in order to make sure they were able to follow the instructions (especially important when no other behavioral measure was taken). For the AT conditions participants were also asked to rate the level of internal speech, internal visualization and interest on a five-point scale (1 = very low to 5 = very high). For the VF they were asked to count the generated words.

### Pre-processing and statistical data Analysis

fMRI Data were analyzed with brain-voyager QX and matlab 2014b with the Neuroelf toolbox version 0.9. The functional images were incorporated into the 3D data sets through trilinear interpolation. Preprocessing of functional scans included 3D motion correction, filtering out of low frequencies (high pass) up to 2 cycles per scan (slow drift), and spatial smoothing using a Gaussian kernel with a full width at half maximum of 6 mm.

Segments showing movement artifacts larger than 1 mm or sharp head movements were excluded from the analysis. The cortical surface in a Talairach coordinate system ^56^ was reconstructed for each participant from the 3D-spoiled gradient echo scan. The functional images were then superimposed on 2D anatomic images and incorporated into the 3D data sets through trilinear interpolation.

Statistical analysis was based on the general linear model (GLM), performed separately for each participant and for each scan with a regressor for each condition, excluding the rest epochs. All regressors were modeled as box-car functions convolved with the hemodynamic response function. In order to account for non-neuronal contributions to the BOLD signal, six null predictors were added to the analysis, corresponding to head motion in the relevant axes (3 translational and 3 rotational vectors). The analysis was performed independently for the time-course of each individual voxel.

The multi-participant functional maps were projected on an inflated or unfolded Talairach normalized brain. Significance levels were calculated, taking into account the minimum cluster size and the probability threshold of a false detection of any given cluster. This was accomplished by a Monte-Carlo simulation (cluster-level statistical threshold estimator in “Brain-voyager”), targeting a threshold of p <.05 corrected (using 1500 iterations). The probability of a false positive detection per image was determined from the frequency count of cluster sizes within the entire brain. For the AT1 v. rest a minimum cluster size of 16 voxels was significant, for AT2 vs. rest - 12 voxels, AT2 vs. Sentences Conjugation – 12 voxels, Imagine Navigation vs. rest – 18 voxels, Imagine Tools vs. rest – 19 voxels, AT1 vs. Verbal Fluency – 17 voxels, Verbal Fluency vs. rest – 25 voxels, Visual Categories vs. rest – 31 voxels, Sentences Conjugation vs. rest – 29 voxels, Working Memory vs. rest – 32 voxels, rest vs. Sentences Conjugation – 28 voxels. For presentation purposes we used the value 25 as cluster threshold for the maps of the first six contrasts, and cluster threshold of 32 voxels for the last four contrasts.

For each participant and a given ROI we extracted the mean time course across all ROI voxels and calculated the event-related averaged response for the conditions in each scan. The signal was normalized to percent signal change relative to a baseline which was defined as the average value of the 2 TRs prior to the stimulus. Additionally, using the results from the GLM, we calculated the mean beta value for each ROI and participant separately. The comparison between the beta values in the two AT scans was done using two-tailed *t*-test, Bonferroni corrected for multiple comparisons.

### Neural pattern classifier

In order to perform neural pattern classification, we conducted “Leave one out” analysis. This was performed for five bulk conditions, lumping together for each participant the average betas of the two thought conditions, the two language conditions, the working memory task, the two imagery conditions and the four visual conditions. Then, in order to avoid highly unreliable voxels, which may introduce noise to the analysis, we performed again the analyses (as previously done by ^57^; and by ^58^), this time including only the 50% of total voxels with lowest variability. We calculated the correlation coefficients per participant and mean group betas (not including the “leave” participant). This allowed us to classify for each participant the task with highest correlation coefficient; correct classifications were considered highest correlation with AT bulk conditions. Percent participants resulting with correct classification was calculated. This was performed for the whole cortex (100% cortical voxels) and 50% low variability voxels.

For the individual tasks classification, in order to calculate the statistical significance, we randomly permuted the tasks in each participant and calculated the correlations coefficients with the mean of the group as described above, this was repeated 10k times. This allowed us to evaluate the probability to achieve such classification by chance, with lower than 5% considered significant.

### Neuronal distance metric

In order to further probe the relationship between abstract thought and other tasks aside from overlap of univariate activity, we added a Neuronal Distance Analyses. In this analyses, a vector was constructed by concatenating the mean beta values for each voxel (across the entire cortex) in each participant and each condition. The neural distance between 2 tasks for the same participant was defined as the Euclidean distance between the corresponding task vectors, similar as in other studies ^57, 59^.

Correlation coefficients were calculated per participant between beta vectors for each pair of conditions to produce the pair-wise distance matrix (higher correlation coefficient – shorter distance). The average distance matrix of all participants is displayed (n=20, figure S1). Average ± SEM correlation coefficients between abstract thought conditions (thought 1 and thought 2) and any other condition were shown in the bar histograms (Figure 8). Paired t-tests were performed between the correlation coefficient of AT1 and AT2 compared to the correlations between thought 1 and the other conditions. (figure 8A, and figure 8B respectively). Regarding the issue of multiple-comparisons, it should be noted that the critical comparison is between the thoughts and the language activations - so basically we have two comparisons (and all the other are given for reference). To minimize this concern, we also acquired five bulk conditions, lumping together for each participant the average correlation coefficient of the two thought conditions, the two language conditions (regarding the language working memory task as a different category), the two imagery conditions and the four visual conditions (figure 8A&B). Average ± SEM correlation coefficients between abstract thought conditions and the other conditions are also shown in the bar histogram (figure 8C).

### Inter-participant variability

In order to estimate the inter-participant variability in the different conditions, for each condition whole brain mean beta vector (cortex only) was calculated and correlation coefficient of each participant to the mean beta was calculated.

### 1.1 Definitions of ROIs

In all the ROI analyses described, ROIs of activation were defined for each participant individually, as the activated voxels located within 10^3 mm around the activation/deactivation peaks, as will be described below. A functional ROI approach in which the region is first identified functionally in each participant individually was chosen because functional properties are more consistently associated with functional ROIs than with locations in stereotaxic space, and is best suited to test functional hypotheses ^60, 61^.

Language ROIs, the L-Broca and L-IFG were defined from the sentence-conjugation task, as the activated voxels located within 10^3 mm of the activity center. DMN ROIs were selected based on resting-state functional connectivity. The seed for the functional-connectivity analysis was the left precuneus (Prc), defined from the Visual Categories scan as the voxels within 10^3 mm around the point of highest deactivation in the respective anatomical region, confirmed using an Atlas ^62^. The other DMN ROIs, right Prc and the bilateral posterior part of inferior parietal lobule (IPL), were identified by lowering the threshold of the functional connectivity map and choosing the highest functionally connected respective regions which appear in specific anatomical locations (thus ensuring the correct selection of DMN regions using both functional and anatomical criteria). The network maps were inspected individually and compared between participants to ensure that the ROIs did not considerably vary anatomically. We found for the L-Prc that the inter-individual differences across participants did not exceed 15 voxels (mean of 6 voxels).

## Availability statement

All the data is publically available at the following link https://drive.google.com/drive/folders/1KzA7Bo1otaEEhBykan1v4l_KTDeNBxL7?usp=sharing

## Code availability

All the custom codes are publically available at the following link https://drive.google.com/drive/folders/1KzA7Bo1otaEEhBykan1v4l_KTDeNBxL7?usp=sharing

## Acknowledgements

The study was funded by the Helen and Kimmel Award for innovative Research, The EU (FP7 VERE), The EU - Human Brain Project and the ISF-ICORE grants to Prof. R. Malach, and the Teva Pharmaceutical Industries LTD fellowship to Dr. A. Berkovich-Ohana. Dr. E. Furman-Haran holds the Calin and Elaine Rovinescu Research Fellow Chair for Brain Research.

