## Supplementary material for "Inter-thinker consistency of language activations during abstract thoughts": Supplemetary figures and tables

Supplementary figures

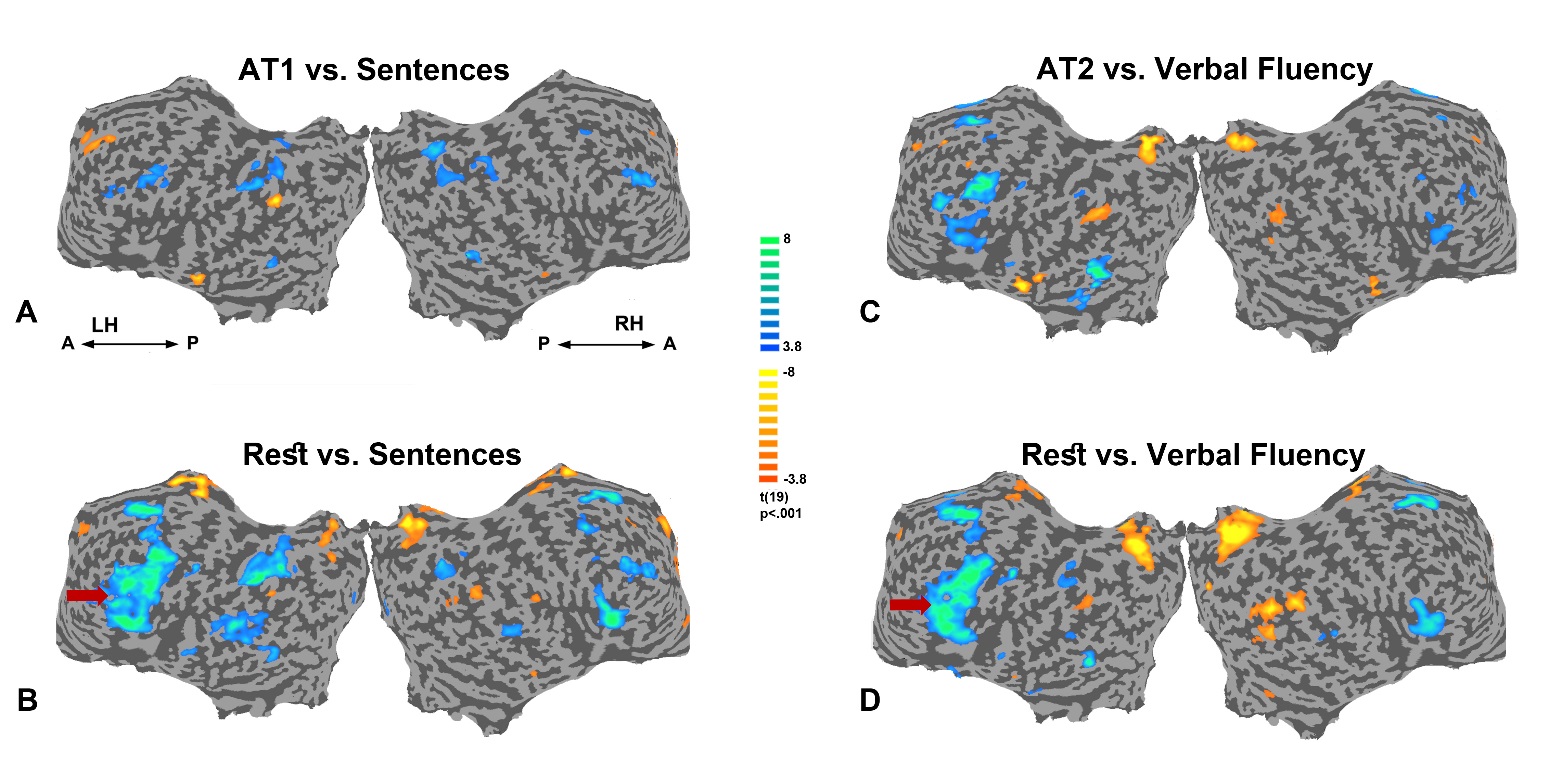

**Figure S1: Activations during the abstract thought and rest relative to language tasks.** Whole brain maps (multi-subject random-effect GLM analysis, *n* = 20, Monte Carlo corrected). A. Abstract-thought 1 relative to sentences conjugation; B. Rest relative to sentences conjugation; C. Abstract-thought 2 relative to Verbal Fluency; E. Rest relative to Verbal Fluency. Color bar indicates activations in yellow, and deactivations in blue. Unfolded views: LH - left hemisphere; RH - right hemisphere; A - anterior; P – posterior.

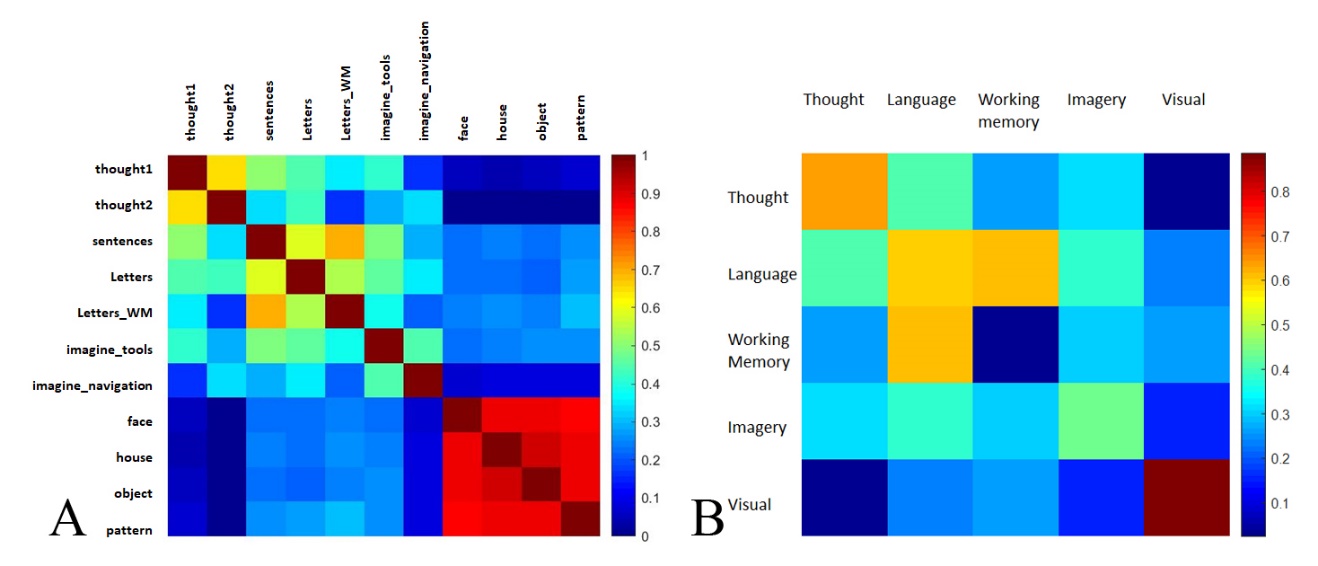

**Figure S2: Neuronal Distance Analyses as correlation coefficient matrices**. A correlation coefficient matrix between the vectors, constructed by concatenating the mean beta for each voxel (in the entire grey matter) in each subject and each condition, and averaged across subjects (*n* = 20), shown **A.** for each pairs of conditions separately; and **B.** between “bulks” of similar conditions (grouped as follows: Thought - AT1 and AT2; Language – sentences and verbal fluency; Imagery – imagine tools and imagine navigation; Visual – objects, houses, patterns, and faces). Notice that shorter distances give warmer colors.

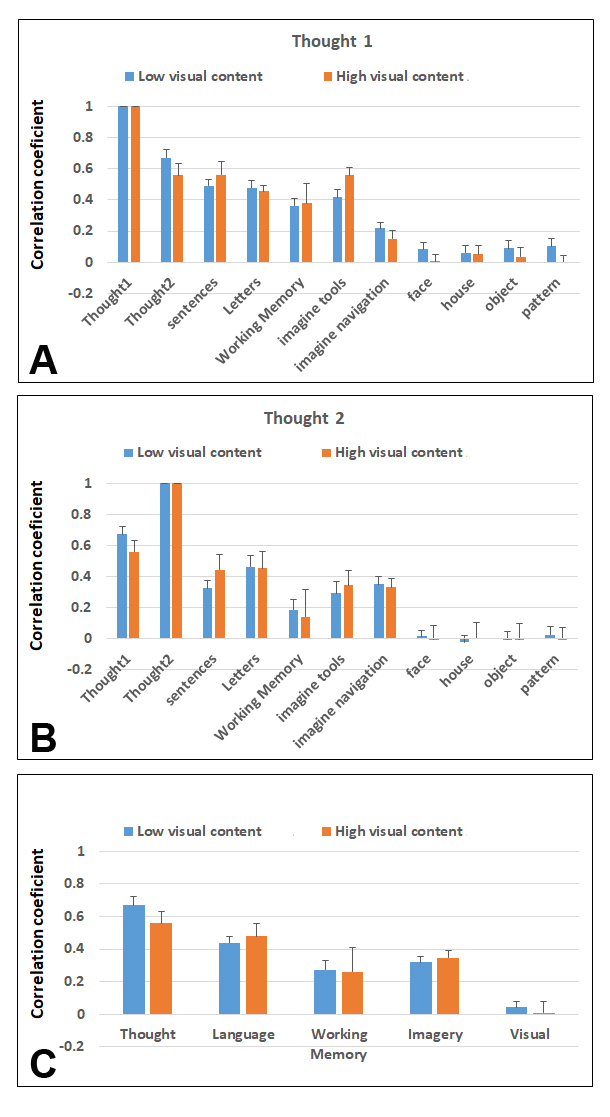

**Figure S3: *Comparing different phenomenal categories.*** *Participants were divided into to two categories, low visual content (n=12, grading visual content as 1 or 2) and high visual content (n=3, grading visual content as 4 or 5). Average (Mean ± SEM) correlation coefficient between AT1 (A), AT2 (B) and bulk AT (C) with other conditions for the two categories.*

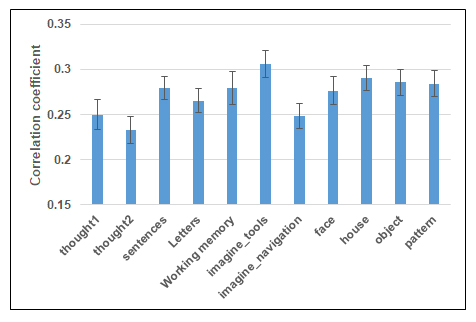

Supplementary tables

**Supplementary table S1: Regions of Interest (ROIs) that exhibited significantly lower or higher activity during abstract Thought 1 (AT1) relative to Rest.** Coordinates are reported in Talairach space. The displayed t-values are associated with the area’s lowest/highest hemodynamic response during AT1 relative to Rest blocks. All coordinates emerged at a threshold of p < 0.05, number of voxels = 16, Monte Carlo corrected. BA, Brodmann area.

| **Grey matter regions** | | **Coordinates** | | | **Statistics** | |
| --- | --- | --- | --- | --- | --- | --- |
| **ROI** | **BA** | **X** | **Y** | **Z** | **T values** | **P values** |
| Right Cerebrum, Parietal lobe, supramarginal gyrus | 40 | 60 | -43 | 37 | -4.320785 | 0.000412 |
| Right Cerebrum, Superior temporal gyrus | 41 | 42 | -37 | 4 | 5.437042 | 0.000036 |
| Right Cerebrum, Superior temporal gyrus | 38 | 45 | 17 | -17 | 5.248569 | 0.000054 |
| Right Cerebrum, Inferior Semi-Lunar Lobule | NA | 24 | -61 | -41 | 7.963513 | 0.000000 |
| Right Cerebrum, Caudate | NA | 15 | 5 | 19 | 5.453264 | 0.000035 |
| Left Cerebrum, Cingulate Gyrus | 31 | -9 | -37 | 40 | -4.820770 | 0.000137 |
| Left Cerebrum, Superior frontal gyrus | 6 | -9 | -1 | 61 | 11.521506 | 0.000000 |
| Left Cerebrum, Putamen | NA | -18 | 2 | 16 | 7.910404 | 0.000000 |
| Left Cerebrum, Inferior frontal gyrus | 9 | -54 | 17 | 22 | 8.212461 | 0.000000 |
| Left Cerebrum, Temporal Lobe, Middle Temporal Gyrus | NA | -63 | -40 | 1 | 7.681249 | 0.000000 |

**Supplementary table S2: Regions of Interest (ROIs) that exhibited significantly lower or higher activity during abstract Thought 2 (AT2) relative to Rest.** Coordinates are reported in Talairach space. The displayed t-values are associated with the area’s lowest/highest hemodynamic response during AT2 relative to Rest blocks. All coordinates emerged at a threshold of p < 0.05, number of voxels = 12, Monte Carlo corrected. BA, Brodmann area.

| **Grey matter regions** | | **Coordinates** | | | **Statistics** | |
| --- | --- | --- | --- | --- | --- | --- |
| **ROI** | **BA** | **X** | **Y** | **Z** | **T values** | **P values** |
| Right Cerebrum, Parietal Lobe, Supramarginal Gyrus | 40 | 63 | -43 | 37 | -5.514978 | 0.000026 |
| Right Cerebellum, Posterior Lobe, Pyramis | - | 21 | -64 | -29 | 5.781610 | 0.000014 |
| Right Cerebellum, Posterior Lobe, Cerebellar Tonsil | - | 3 | -49 | -35 | 6.041409 | 0.000008 |
| Left Cerebrum, Frontal Lobe, Superior Frontal Gyrus | 6 | -9 | 5 | 67 | 6.781460 | 0.000002 |
| Left Cerebrum, Sub-lobar, Caudate | - | -15 | 5 | 13 | 5.264611 | 0.000044 |
| Left Cerebrum, Parietal Lobe, Postcentral Gyrus | 3 | -54 | -10 | 52 | 5.805932 | 0.000014 |
| Left Cerebrum, Frontal Lobe, Inferior Frontal Gyrus | 45 | -48 | 26 | 4 | 10.243520 | 0.000000 |
| Left Cerebrum, Temporal Lobe, Middle Temporal Gyrus | 22 | -48 | -37 | 1 | 5.215112 | 0.000049 |
| Left Cerebrum, Temporal Lobe, Middle Temporal Gyrus | 21 | -63 | -10 | -11 | 5.436362 | 0.000030 |

**Supplementary table S3: Regions of Interest (ROIs) that exhibited significantly lower or higher activity during Imagine Navigation relative to Rest.** Coordinates are reported in Talairach space. The displayed t-values are associated with the area’s lowest/highest hemodynamic response during Imagine Navigation relative to Rest blocks. All coordinates emerged at a threshold of p < 0.05, number of voxels = 18, Monte Carlo corrected. BA, Brodmann area.

| **Grey matter regions** | | **Coordinates** | | | **Statistics** | |
| --- | --- | --- | --- | --- | --- | --- |
| **ROI** | **BA** | **X** | **Y** | **Z** | **T values** | **P values** |
| Right Cerebrum, Limbic Lobe, Posterior Cingulate | 30 | 12 | -55 | 13 | 12.524324 | 0.000000 |
| Right Cerebrum, Sub-lobar, Insula | 13 | 45 | -13 | 13 | -4.665721 | 0.000168 |
| Right Cerebrum, Frontal Lobe, Middle Frontal Gyrus | 9 | 36 | 11 | 25 | 4.814264 | 0.000121 |
| Right Cerebrum, Limbic Lobe, Parahippocampal Gyrus | 36 | 27 | -34 | -14 | 10.469896 | 0.000000 |
| Right Cerebellum, Anterior Lobe, Culmen | - | 9 | -67 | -8 | -4.681157 | 0.000163 |
| Right Cerebrum, Frontal Lobe, Sub-Gyral | 6 | 27 | 2 | 52 | 9.452079 | 0.000000 |
| Right Cerebrum, Sub-lobar, Claustrum | - | 27 | 20 | 10 | 5.080540 | 0.000066 |
| Left Cerebrum, Limbic Lobe, Cingulate Gyrus | 31 | -3 | -28 | 37 | -5.077569 | 0.000067 |
| Left Cerebrum, Occipital Lobe, Cuneus | 18 | -18 | -91 | 19 | -5.745980 | 0.000015 |
| Left Cerebrum, Sub-lobar, Caudate | - | -15 | 2 | 13 | 4.333264 | 0.000358 |
| Left Cerebellum, Posterior Lobe, Cerebellar Tonsil | - | -42 | -40 | -38 | 6.512733 | 0.000003 |
| Left Cerebrum, Sub-lobar, Insula | 13 | -30 | 20 | 7 | 5.734355 | 0.000016 |
| Left Cerebrum, Parietal Lobe, Inferior Parietal Lobule | 40 | -33 | -49 | 37 | 6.119219 | 0.000007 |

**Supplementary table S4: Regions of Interest (ROIs) that exhibited significantly lower or higher activity during Imagine Tools relative to Rest.** Coordinates are reported in Talairach space. The displayed t-values are associated with the area’s lowest/highest hemodynamic response during Imagine Tools relative to Rest blocks. All coordinates emerged at a threshold of p < 0.05, number of voxels = 19, Monte Carlo corrected. BA, Brodmann area.

| **Grey matter regions** | | **Coordinates** | | | **Statistics** | |
| --- | --- | --- | --- | --- | --- | --- |
| **ROI** | **BA** | **X** | **Y** | **Z** | **T values** | **P values** |
| Right Cerebrum, Temporal Lobe, Middle Temporal Gyrus | 21 | 57 | 8 | -20 | -7.679189 | 0.000000 |
| Right Cerebrum, Parietal Lobe, Postcentral Gyrus | 43 | 51 | -10 | 16 | -5.386225 | 0.000041 |
| Right Cerebrum, Sub-lobar, Insula | 13 | 45 | -25 | 25 | -4.436769 | 0.000318 |
| Right Cerebrum, Frontal Lobe, Middle Frontal Gyrus | 47 | 51 | 38 | -5 | -6.133943 | 0.000009 |
| Right Cerebrum, Frontal Lobe, Superior Frontal Gyrus | 8 | 42 | 20 | 46 | -6.371567 | 0.000005 |
| Right Cerebellum, Anterior Lobe, Culmen | - | 30 | -55 | -26 | 4.593233 | 0.000226 |
| Right Cerebrum, Frontal Lobe, Medial Frontal Gyrus | 8 | 6 | 50 | 40 | -7.104651 | 0.000001 |
| Right Cerebrum, Limbic Lobe, Cingulate Gyrus | 31 | 9 | -34 | 40 | -7.380973 | 0.000001 |
| Left Cerebrum, Frontal Lobe, Medial Frontal Gyrus | 6 | -12 | 2 | 55 | 5.313161 | 0.000047 |
| Left Cerebrum, Limbic Lobe, Posterior Cingulate | 29 | -15 | -49 | 10 | 6.315782 | 0.000006 |
| Left Cerebrum, Parietal Lobe, Superior Parietal Lobule | 7 | -27 | -70 | 46 | 6.762531 | 0.000002 |
| Left Cerebrum, Frontal Lobe, Sub-Gyral | 6 | -27 | -4 | 55 | 5.620329 | 0.000025 |
| Left Cerebrum, Temporal Lobe, Fusiform Gyrus | 37 | -27 | -37 | -14 | 7.800393 | 0.000000 |
| Left Cerebellum, Posterior Lobe, Inferior Semi-Lunar Lobule | - | -24 | -73 | -35 | -4.818781 | 0.000138 |

**Supplementary table S5: Regions of Interest (ROIs) that exhibited significantly lower or higher activity during Verbal Fluency relative to Rest.** Coordinates are reported in Talairach space. The displayed t-values are associated with the area’s lowest/highest hemodynamic response during Verbal Fluency relative to Rest blocks. All coordinates emerged at a threshold of p < 0.05, number of voxels = 25, Monte Carlo corrected. BA, Brodmann area.

| **Grey matter regions** | | **Coordinates** | | | **Statistics** | |
| --- | --- | --- | --- | --- | --- | --- |
| **ROI** | **BA** | **X** | **Y** | **Z** | **T values** | **P values** |
| Right Cerebrum, Temporal Lobe, Superior Temporal Gyrus | 22 | 54 | -55 | 19 | -7.583309 | 0.000000 |
| Left Cerebrum, Frontal Lobe, Inferior Frontal Gyrus | 9 | -51 | 8 | 31 | 11.867943 | 0.000000 |
| Right Cerebrum, Temporal Lobe, Superior Temporal Gyrus | 38 | 51 | 17 | -26 | -4.343438 | 0.000350 |
| Right Cerebrum, Temporal Lobe, Superior Temporal Gyrus | 22 | 54 | -25 | 1 | 5.344560 | 0.000037 |
| Right Cerebellum, Anterior Lobe, Culmen | - | 33 | -58 | -26 | 6.873565 | 0.000001 |
| Right Cerebrum, Limbic Lobe, Cingulate Gyrus | 31 | 6 | -58 | 28 | -10.399951 | 0.000000 |
| Right Cerebrum, Frontal Lobe, Superior Frontal Gyrus | 9 | 12 | 53 | 31 | -4.588031 | 0.000201 |
| Left Cerebrum, Limbic Lobe, Anterior Cingulate | 24 | 0 | 32 | 10 | -5.151993 | 0.000057 |
| Left Cerebellum, Posterior Lobe, Declive | - | -18 | -73 | -11 | -5.046850 | 0.000072 |
| Left Cerebrum, Parietal Lobe, Inferior Parietal Lobule | 40 | -42 | -49 | 43 | 5.301098 | 0.000041 |
| Left Cerebrum, Temporal Lobe, Fusiform Gyrus | 37 | -48 | -52 | -11 | 8.379123 | 0.000000 |

**Supplementary table S6: Regions of Interest (ROIs) that exhibited significantly lower or higher activity during Sentences Conjugation relative to Rest.** Coordinates are reported in Talairach space. The displayed t-values are associated with the area’s lowest/highest hemodynamic response during Sentences Conjugation relative to Rest blocks. All coordinates emerged at a threshold of p < 0.05, number of voxels = 12, Monte Carlo corrected. BA, Brodmann area.

| **Grey matter regions** | | **Coordinates** | | | **Statistics** | |
| --- | --- | --- | --- | --- | --- | --- |
| **ROI** | **BA** | **X** | **Y** | **Z** | **T values** | **P values** |
| Left Cerebrum, Frontal Lobe, Precentral Gyrus | 6 | -36 | -1 | 40 | 11.301556 | 0.000000 |
| Right Cerebrum, Parietal Lobe, Supramarginal Gyrus | 40 | 51 | -52 | 31 | -5.569594 | 0.000028 |
| Right Cerebrum, Parietal Lobe, Inferior Parietal Lobule | 40 | 48 | -28 | 25 | -5.229927 | 0.000057 |
| Right Cerebrum, Temporal Lobe, Superior Temporal Gyrus | 22 | 57 | -13 | 4 | 5.818115 | 0.000016 |
| Right Cerebrum, Temporal Lobe, Middle Temporal Gyrus | 21 | 51 | -1 | -23 | -4.978267 | 0.000097 |
| Right Cerebellum, Anterior Lobe | NA | 18 | -64 | -26 | 8.903316 | 0.000000 |
| Right Cerebrum, Parietal Lobe, Precuneus, | 7 | 9 | -73 | 52 | 4.665727 | 0.000192 |
| Right Cerebrum, Frontal Lobe, Superior Frontal Gyrus | 8 | 15 | 38 | 52 | -5.008143 | 0.000091 |
| Right Cerebrum, Frontal Lobe, Medial Frontal Gyrus | 9 | 3 | 47 | 16 | -6.216230 | 0.000007 |
| Left Cerebrum, Limbic Lobe, Cingulate Gyrus | 24 | 0 | -22 | 37 | -8.330937 | 0.000000 |
| Right Cerebrum, Occipital Lobe, Lingual Gyrus | 18 | 3 | -85 | 1 | 4.616590 | 0.000214 |
| Left Cerebrum, Temporal Lobe, Superior Temporal Gyrus | 39 | -48 | -58 | 28 | -5.088785 | 0.000077 |

**Supplementary table S7: Regions of Interest (ROIs) that exhibited significantly lower or higher activity during Working Memory relative to Rest.** Coordinates are reported in Talairach space. The displayed t-values are associated with the area’s lowest/highest hemodynamic response during Working Memory relative to Rest blocks. All coordinates emerged at a threshold of p < 0.05, number of voxels = 32, Monte Carlo corrected. BA, Brodmann area.

| **Grey matter regions** | | **Coordinates** | | | **Statistics** | |
| --- | --- | --- | --- | --- | --- | --- |
| **ROI** | **BA** | **X** | **Y** | **Z** | **T values** | **P values** |
| Right Cerebrum, Temporal Lobe, Superior Temporal Gyrus | 22 | 66 | -61 | 16 | -5.047945 | 0.000084 |
| Left Cerebrum, Parietal Lobe, Inferior Parietal Lobule | 40 | -42 | -46 | 40 | 13.449508 | 0.000000 |
| Right Cerebrum, Temporal Lobe, Middle Temporal Gyrus | 37 | 51 | -64 | 4 | -3.417123 | 0.003073 |
| Right Cerebrum, Frontal Lobe, Superior Frontal Gyrus | 11 | 21 | 62 | -26 | 4.452213 | 0.000308 |
| Right Cerebrum, Parietal Lobe, Precuneus | 31 | 6 | -61 | 25 | -8.310966 | 0.000000 |
| Left Cerebrum, Limbic Lobe, Anterior Cingulate, | 24 | 0 | 26 | -2 | -5.372716 | 0.000042 |
| Left Cerebrum, Temporal Lobe, Middle Temporal Gyrus | 39 | -54 | -79 | 25 | -5.871235 | 0.000015 |

**Supplementary table S8: Regions of Interest (ROIs) that exhibited significantly lower or higher activity during Visual Categories relative to Rest.** Coordinates are reported in Talairach space. The displayed t-values are associated with the area’s lowest/highest hemodynamic response during Visual Categories relative to Rest blocks. All coordinates emerged at a threshold of p < 0.05, number of voxels = 31, Monte Carlo corrected. BA, Brodmann area.

| **Grey matter regions** | | **Coordinates** | | | **Statistics** | |
| --- | --- | --- | --- | --- | --- | --- |
| **ROI** | **BA** | **X** | **Y** | **Z** | **T values** | **P values** |
| Right Cerebrum, Temporal Lobe, Middle Temporal Gyrus | 21 | 45 | 2 | -23 | -8.728031 | 0.000000 |
| Right Cerebrum, Temporal Lobe, Middle Temporal Gyrus | 39 | 48 | -64 | 28 | -7.039969 | 0.000001 |
| Right Cerebrum, Occipital Lobe, Middle Occipital Gyrus | 18 | 30 | -85 | -2 | 15.622702 | 0.000000 |
| Left Cerebrum, Occipital Lobe, Lingual Gyrus | 18 | -24 | -94 | -5 | 12.634987 | 0.000000 |
| Right Cerebrum, Frontal Lobe, Inferior Frontal Gyrus | 45 | 57 | 38 | 4 | -5.274949 | 0.000051 |
| Left Cerebrum, Frontal Lobe, Superior Frontal Gyrus | 8 | -21 | 20 | 49 | -10.934224 | 0.000000 |
| Left Cerebrum, Limbic Lobe, Posterior Cingulat | 31 | -6 | -55 | 22 | -12.179070 | 0.000000 |
| Right Cerebrum, Limbic Lobe, Parahippocampal Gyrus, Hippocampus | NA | 27 | -22 | -11 | -4.504663 | 0.000274 |
| Right Cerebrum, Sub-lobar, Thalamus, Gray Matter, Ventral Anterior Nucleu | NA | 12 | -4 | 10 | 4.407712 | 0.000340 |
| Left Cerebellum, Posterior Lobe, Inferior Semi-Lunar Lobule | NA | -6 | -70 | -38 | 5.314754 | 0.000047 |
| Right Cerebrum, Occipital Lobe, Cuneus | 18 | 6 | -97 | 22 | -3.978165 | 0.000882 |
| Left Cerebrum, Limbic Lobe, Parahippocampal Gyrus | 28 | -24 | -16 | -20 | -4.695185 | 0.000180 |
| Left Cerebrum, Sub-lobar, Insula | 13 | -30 | 20 | 10 | 6.285265 | 0.000006 |
| Left Cerebrum, Temporal Lobe, Middle Temporal Gyrus | 21 | -67 | -13 | -8 | -5.581305 | 0.000027 |
| Left Cerebrum, Temporal Lobe, Middle Temporal Gyrus | 39 | -48 | -64 | 22 | -8.594175 | 0.000000 |
| Left Cerebrum, Frontal Lobe, Inferior Frontal Gyrus | 45 | -54 | 35 | 1 | -6.683321 | 0.000003 |
